# Evolutionary Perspective and Expression Analysis of Intronless Genes Highlight the Conservation on Their Regulatory Role

**DOI:** 10.1101/2021.01.13.426573

**Authors:** Katia Aviña-Padilla, José Antonio Ramírez-Rafael, Gabriel Emilio Herrera-Oropeza, Vijaykumar Muley, Dulce I. Valdivia, Erik Díaz-Valenzuela, Andrés García-García, Alfredo Varela-Echavarría, Maribel Hernández-Rosales

**Affiliations:** Instituto de Neurobiología, Universidad Nacional Autónoma de México, Querétaro, México; Centro de Investigación y de Estudios Avanzados del IPN, Unidad Irapuato, Guanajuato, México; Centro de Física Aplicada y Tecnología Avanzada, Universidad Nacional Autónoma de México, Querétaro, México; Centre for Developmental Neurobiology, Institute of Psychiatry, Psychology, and Neuroscience, King’s College London, London, United Kingdom

**Keywords:** intronless genes_1_, exon-intron architecture_2_, embryonic telencephalon_3_, protocadherins_4_, histones_5_, transcription factors_6_, evolutionary histories_7_, microproteins_8_

## Abstract

Eukaryotic gene structure is a combination of exons generally interrupted by intragenic non-coding DNA regions termed introns removed by RNA splicing to generate the mature mRNA. Thus, eukaryotic genes can be either single exon genes (SEGs) or multiple exon genes (MEGs). Among SEGs, intronless genes (IGs) are a subgroup that additionally lacks introns at their UTRs, and code for proteins essentially involved in development, growth, and cell proliferation. Gene expression of IGs has been proposed to be highly specialized for neuro-specific functions and linked to cancer, neuropathies, and developmental disorders. The abundant presence of introns in eukaryotic genomes is pivotal for the precise control of gene expression. Notwithstanding, IGs exempting splicing events entail a higher transcriptional fidelity, making them even more valuable for regulatory roles. This work aimed to infer the functional role and evolutionary history of IGs using the mouse genome. Intronless protein-coding genes consist of a subgroup of ~6 % of a total of 21,527 genes with one exon. To understand the prevalence, biological relevance, and evolution, we identified and studied their 1,116 functional proteins. We validated differential expression in transcriptomics data of early embryo stages using mouse telencephalon tissue. Our results showed that expression levels of IGs are lower compared to MEGs. However, strongly upregulated IGs include transcription factors (TFs) such as the class 3 of POU (HMG Box), *Neurog1, Olig1*, and *BHLHe22, BHLHe23,* among other essential genes including the beta cluster of protocadherins. Most striking was the finding that IG-encoded BHLH TFs qualify the criteria to be referred to as microprotein candidates. Finally, predicted protein orthologs in other six genomes confirmed a high conservancy of IGs associated with regulating neurobiological processes and with chromatin organization and epigenetic regulation in *Vertebrata*. Moreover, this study highlights that IGs are essential modulators of regulatory processes, as Wnt signaling pathway and biological processes as pivotal as sensory organs developing at a transcriptional and post-translational level. Overall, our results suggest that IG proteins have specialized, prevalent, and unique biological roles and that functional divergence between IGs and MEGs is likely to be the result of specific evolutionary constraints.

## Introduction

Genes with one exon are characteristic of prokaryotic genomes. Most eukaryotic genes, in contrast, contain introns, nucleotide DNA sequences that after transcription as part of the messenger RNA are removed by splicing during its maturation. The latter are thus termed multiple exon genes (MEGs), although eukaryotic genomes also contain an important proportion of single-exon genes (SEGs).

Diverse studies of SEGs in eukaryotes have been performed over the past decades (Tine et al. 2011; Zou, Guo, and He 2011; K. R. Sakharkar et al. 2006; Yan et al. 2014). A precise definition, however, has only been recently formulated. The improved ontology defines an SEG as a nuclear gene whose protein-coding sequence comprises only one exon. Pseudogenes, functional RNAs, tRNA, rRNA, ribozyme long non-coding RNAs, and miRNAs are excluded from this definition. This classification scheme includes two main groups of SEGs; those having introns in their untranslated regions ‘uiSEGs’; and those lacking introns in the entire gene, termed ‘Intronless Genes’ (IGs) (Jorquera et al. 2018).

Because of their prokaryotic-like architecture, IGs in eukaryotic genomes, provide interesting datasets for computational analysis in comparative genomics and evolutionary trajectories. Comparative analysis of their sequences between different genomes could help to identify their unique and conserved features, thus providing insights into the role of introns in gene evolution and lead to a better understanding of genome architecture and arrangement.

The abundant presence of introns in most genes of multicellular organisms entails regulatory processes associated with the generation of multiple splice variants not present in intronless genes. In consequence, IGs not requiring splicing events represents a higher transcriptional fidelity, which makes them even more valuable for regulatory roles. To date, more than 2000 protein-coding genes in the human genome have been determined as SEGs (https://www.ensembl.org/index.html). Among them, a considerable amount of IGs is responsible for encoding proteins such as G-protein-coupled receptors (GPCRs), canonical histones, transcription factors, many genes involved in signal transduction, as well as some involved in the regulation of development, growth, and cell proliferation (Meena K Sakharkar et al. 2005; Grzybowska 2012).

The expression of IGs has been proposed to be highly specialized for neuro-specific functions and linked to diseases such as cancer, neuropathies, and developmental disorders. Examples of IGs with clinical relevance are the *RPRM* gene related to gastric cancer which causes increased cell proliferation and possesses tumor suppression activity (Amigo et al. 2018) and the protein kinase *CK2α* gene which is up-regulated in all human cancers (Hung et al. 2010). Other IGs linked to cancer include CLDN8 in colorectal carcinoma and renal cell tumors, *ARLTS1* in melanoma, and *PURA* and *TAL2* in leukemia (Grzybowska 2012). IGs have also been associated with neuropathies, such as *ECDR1*, a cerebellar degeneration-related protein, and *NPBWR2*, a neuropeptide B/W receptor type (Louhichi, Fourati, and Rebaï 2011).

Regarding their role in the diseases described above, IGs in humans are potential clinical biomarkers and drug targets that deserve careful consideration (Grzybowska 2012; Ohki et al. 2000; Liu et al. 2017). Their functional role and their evolutionary conservation in other genomes, however, remains poorly understood. Furthermore, there is a current debate between related theories that place the origin of introns, early or late during the evolutionary history of eukaryotes (Fedorova and Fedorov 2003).

With this backdrop, our work aimed to characterize the functional role of mouse IGs and to infer their evolutionary pattern across six additional vertebrate genomes. We have analyzed their expression, particularly during brain development at early embryonic stages, and their potential as transcriptional as well as post-translational modulators.

Overall, this study sheds light on the concerted role played by this peculiar group of genes and helps contrast the functional features of intron-containing and intronless genes across vertebrate species and their collective evolutionary roadmaps.

## Materials and methods

### Data extraction and curation for IG, uiSEG, and MEG datasets

Data were extracted using Python scripts (https://github.com/GEmilioHO/intronless_genes), and the Application Programming Interface (API). Seven vertebrate genomes including *Mus musculus, Homo sapiens, Pan troglodytes, Monodelphis domestica, Rattus norvegicus, Gallus gallus,* and *Danio rerio* assembled at a chromosome level were accessed from the Ensembl REST API platform (http://rest.ensembl.org/ accessed using Python with the ensembl_rest package). The pipeline process was as follows: protein-coding genes with CDS identifiers for transcripts for all chromosomes were retrieved and classified into two datasets named “single-exon genes’’ (SEGs), and “multiple exon genes’’ (MEGs) depending on exon and transcript count (**Suppl. Figure 1**). The first, containing SEGs, was submitted to the Intron DB comparison (http://www.nextgenbioinformatics.org/IntronDB) to separate those with UTR introns and referred to as uiSEGs. The output of the pipeline was a third dataset containing only intronless genes (IGs). After data extraction, a manual curation step in the IG and uiSEG datasets was followed to discard incomplete annotated protein sequences, and mitochondrial encoded proteins (**Suppl. Figure 1**).

### Computational prediction of mouse intronless genes function

Mouse IG and MEG datasets obtained as described before from the Ensembl database according to their exon count were classified into two groups: 1) 20,694 protein-coding genes with more or equal to two exons were labeled as MEGs, 2) 1,116 protein-coding genes with only one exon count and one transcript count were classified as IGs. These were used to perform an over-representation analysis of functional assignment using the SUPERFAMILY (http://supfam.org/; proteins of known three-dimensional structure); Pfam (https://pfam.xfam.org/; protein domains), and PROSITE (https://prosite.expasy.org/; biologically meaningful signatures or motifs) databases. All tests were compared with parallel analysis performed with MEG IDs as controls. For data visualization ClusterProfiler package (G. Yu et al. 2012) in R for SUPERFAMILY and Pfam and Python scripts using a hypergeometric test for PROSITE enrichment was employed.

### Functional Enrichment Analysis of proteins from IG and MEG datasets

The functional enrichment analyses were performed using the Metascape tool (http://metascape.org/), for the biological processes category, including KEGG and Reactome pathways. First, the functional enrichment of the mouse IG proteins was performed. Then, the meta-analysis workflow was used to compare enriched terms for the IG orthologs identified for the 1,116 mouse proteins conserved as IGs in the aforementioned genomes, to the pathways of three random samples of the same size of multi-exon genes. Finally, to determine the conservation of the IGs functional role, we first obtained the overrepresented GO terms for biological processes and molecular function belonging to all the orthologs from the seven genomes using AmiGO2 (http://amigo.geneontology.org/amigo/landing). Then, GO terms with their corresponding p-values were clusterized using REVIGO (Supek et al. 2011), which finds a representative subset of the terms using an algorithm that relies on semantic similarity measures. For data visualization, the Treemap R package was employed (https://www.r-project.org/).

### Data source and differential expression analysis

Read counts from a previous transcriptomic analysis of mouse embryonic telencephalon (E.9.5 and E.10.5) were used to identify differentially expressed genes. The transcriptomes were obtained using the Illumina HiSeq RNA sequencing (RNA-seq) platform. The procedure for read-counts normalization, and to calculate differential expression analysis is described in (https://data.mendeley.com/datasets/rdt5757cbw/1). Mouse IG and MEG datasets were submitted to analysis to determine the directionality of the change in expression at developmental stage B compared to stage A. Genes having significant p-values with positive log2 -fold change represent an increased expression (UP), those with negative log2-fold change are considered with decreased expression (DN), while gene expression with p-values above 0.05 represents no change between stages (NC), and read-count lower than five in less than four samples out of eight are considered not expressed (NE).

### Functional Enrichment Analysis of Differential Expressed IGs and MEGs

The functional enrichment analyses were performed using the over-representation analysis of the functional assignment. Genes with differential expression up to two log2-fold change values were considered as upregulated with a p-value and q-value set at 0.05 and 0.10, respectively. For data analysis, and their visualization the ClusterProfiler package (G. Yu et al. 2012) in R language was employed.

### Post-translational modifications and regulatory assignment of IG proteins

For post-translational modification assignments of IG and MEG proteins, the dbPTM (http://dbptm.mbc.nctu.edu.tw/) was used. A two proportion Z-test was used to check whether the proportions of each post-translational modification among IG and MEG proteins were similar. The p-value was set at 0.05. When the resulting p-value was not significant, meaning that the proportions of IG and MEG proteins were similar for a specific post-translational modification, this was classified as “similar”. On the other hand, when the resulting p-value was < 0.05 the post-translational modification was classified as more abundant in “IG” or “MEG” depending on which one had a higher relative percentage of such modification. Post-translational modifications exclusive of either IG or MEG proteins were classified as “unique”. Then, using the miPFinder program (https://github.com/DaStraub/miPFinder), we determined the mouse gene candidates for IG-encoding microproteins.

### Search for orthologs of mouse IGs

Mouse peptide sequences were submitted to ProteinOrtho (Lechner et al. 2011; 2014) to infer orthologous genes in the genomes of rat, human, chimp, opossum, chicken, and zebrafish. As a first step, ProteinOrtho performs sequence comparison between each pair of genomes and reports best bidirectional hits (BBHs) for alignments with equal or above fifty percent of sequence identity. As a second step, it represents each gene or protein as a node of a graph and places an edge between two genes if they were identified as a BBH, then it applies a clustering algorithm and finally reports the orthology relations as pairs of genes in two different genomes.

Each of the species used in this study is a model organism of a different taxonomic level, and therefore, the conservation of mouse orthologs in close or far related species would give an idea of the “age” of the gene. Therefore, for orthologous genes that were identified in rat, we say that they are conserved in *Muridae*, for those in human and chimp we say they are conserved in *Hominidae*, for those found in opossum, we say they are conserved in *Didelphidae*, for chick, they are conserved in *Phasianidae*, and finally, for those found in zebrafish, they are conserved in *Cyprinidae*. We also identified orthologous genes that were found in *Arabidopsis thaliana* and in *Caenorhabditis elegans* in order to have an idea of those genes that are conserved in plants and in invertebrates, respectively.

### Reconstruction of the evolutionary history of mouse IGs and their conservation in other organisms

From the ProteinOrtho predictions, orthology graphs were constructed, and an in-house developed method for the evolutionary reconstruction of gene families is used. This method implements the theory reported in (Hellmuth et al. 2013; Hernandez-Rosales et al. 2012), and can be found at https://gitlab.com/jarr.tecn/revolutionh-tl. This tool starts by performing a modular decomposition (Tedder et al. 2008) on orthology graphs and then inferring the corresponding gene trees. Each internal node of these trees represents an evolutionary event, duplication, or speciation. Subsequently, the gene trees are reconciled with the species tree to determine in which branch of the species tree events occur and, at the same time, infer gene losses. This method allows us to infer how ancestral a gene is, determined by its orthologs in the other species, as well as to identify species-specific genes. Finally, we identified the orthologs of the 1,116 mouse IG proteins that were conserved as IGs, uiSEGs, or MEGs in the abovementioned genomes.

### Syntenic conservation across β-protocadherin cluster

To determine the syntenic conservation of the mouse protocadherin IG members of the beta cluster, across the selected genomes, the genomic coordinates of orthologs genes were retrieved from GTF files employing custom R scripts and plotted using the genoPlotR R package.

## Results

### Functional Assignment of protein-coding IGs in the Mouse Genome

Computational analysis was performed to identify the functional features and the distribution of IGs across the mouse genome. For the functional assignment, mouse protein-coding IGs representing 6% of the total of genes with one exon in the mouse genome were then analyzed based on the comparative annotation of IG and MEG datasets to identify unique and shared molecular and biological features (**Suppl. Figure 2**).

The grouping of IGs by protein domains that have an evolutionary relationship (SUPERFAMILY database) revealed a higher enrichment of the histone fold, 4−helical cytokine family of signal transducers, and transcription factor families including the POZ domain, “Winged helix” DNA-binding domain, HMG-Box, and A DNA−binding domain in eukaryotic, as well as transmembrane protein families Cadherin−like, and Frizzled cysteine−rich domain when compared to MEGs proteins. Meanwhile, the MEG encoded proteins are enriched in protein kinase-like, immunoglobulin, KRAB domain, and ARM repeat families. The top enriched structural families of IG and MEG groups are shown in **Suppl. Figure 3**.

The analysis of the conserved functional domains (Pfam database) among the enriched protein families encoded by mouse IGs, revealed 598 hits with enriched domains. The molecular function of these proteins was then used to group IGs in the following classes: 253 were transmembrane protein receptors, 101 core histones, 84 transcription factors, and 160 that did not belong to these classes (others) (**Figure 1a**). Among the transmembrane protein receptors, the most enriched domains were TAS2R, V1R, and 7tm_1, common in GPCR and vomeronasal receptors (**Figure 1b**). Other domains identified were cadherin, PMP22_claudin, ADAM CR, Disintegrin, and FZ. In the transcription factor group, BTB, Myb DNA−binding, HMG-Box, forkhead were enriched protein domains in the mouse IGs compared to MEGs (**Figure 1c**). Meanwhile, in the histone group, four domains were observed: Histone, Histone H2AC, CENPTC, and Linker histone (**Figure 1d**). Finally, in the other groups we found among others Keratin, Interferon, Ubiquitin, Actin, FYTT enriched domains (**Figure 1e**). The abovementioned classification of IGs in functional groups was then used for the transcriptional analysis (**Figure 1**).

**Figure 1.**
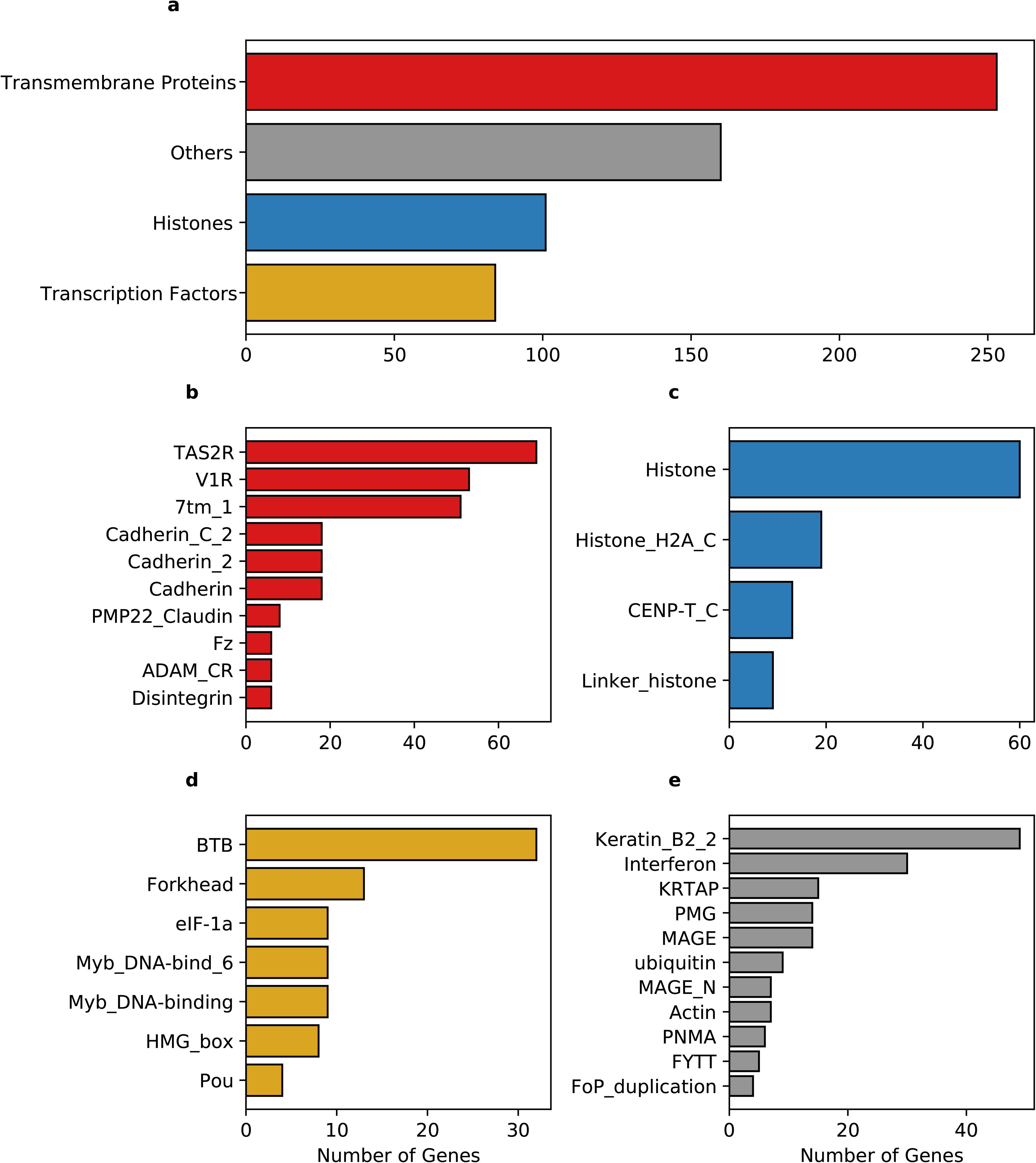
Enrichment of the Pfam protein domains in mouse IG proteins compared to MEG. **a)** Significantly enriched domains of proteins grouped into three main classes, transmembrane proteins (red), histones (blue), and transcription factors (yellow). Enrichment of domain terms is calculated with a significant background list associated with the Pfam domain, and p-adjusted values above 0.05. **b)** Pfam domains grouped in the *transmembrane proteins* class, **c)** Pfam domains grouped in the *histone* class, **d)** Pfam domains grouped in the *transcription factors* class, **e)** Pfam domains considered as *others*.

Analysis of biologically significant motifs (PROSITE database) among MEG and IG proteins identified a total of 1,239 (12,556 hits) and 144 (634 hits) distinct protein signatures, respectively. Interestingly, among the most abundant motifs in the mouse IGs, GPCR, leucine-rich repeat, histone, and transcription factor Forkhead domain and Myc-type bhlh motifs are ankyrins, and cadherin domains more infrequent in MEG proteins (**Figure 2a**). It is noteworthy, however, that among the top motifs that were unique to IG proteins H2B signature (10 hits/10 genes), was the largest group (**Figure 2b**). Hence, these results show that most of the top predictions of IGs signatures are characteristic of transmembrane receptors, histones, and specific transcription factors, having a unique signature for histone.

**Figure 2.**
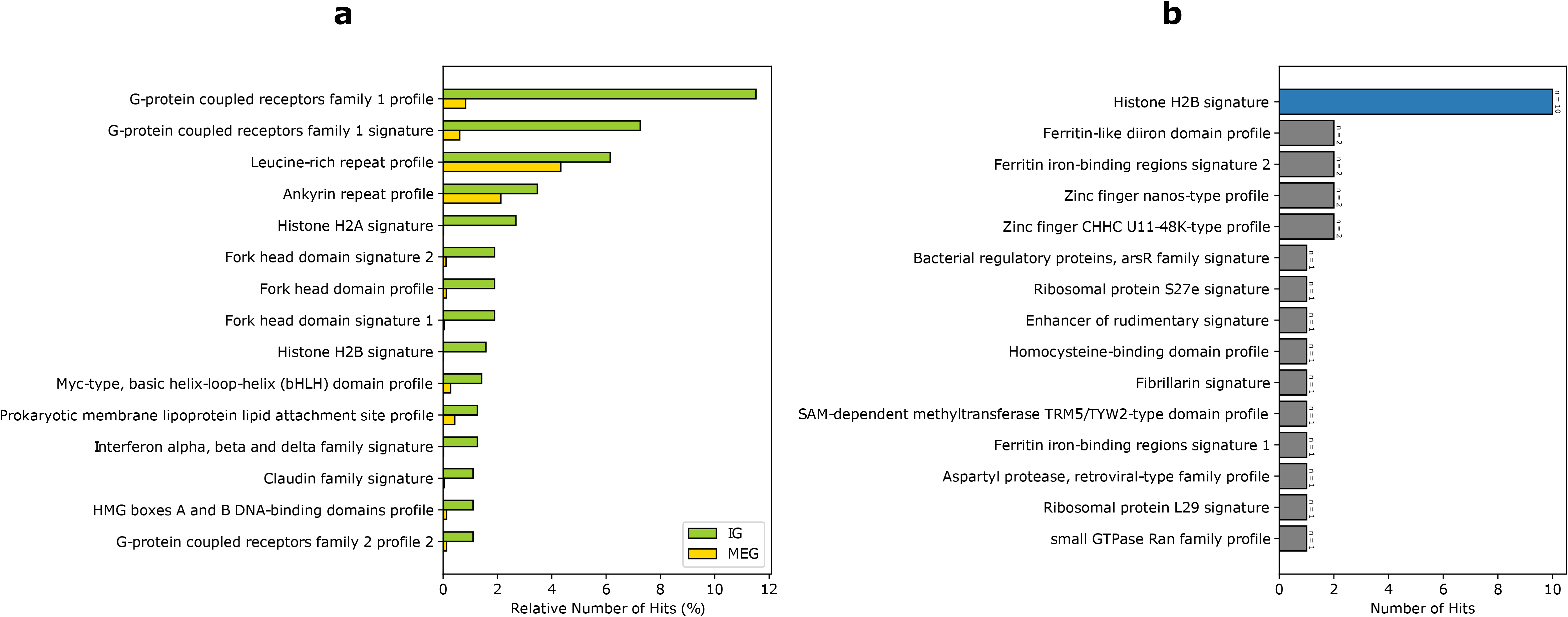
Enrichment of the PROSITE signatures in mouse IG proteins compared to MEG. **a)** 15 most abundant PROSITE signatures of IG proteins. **b)** PROSITE signatures exclusive of IG proteins (signatures belonging to the *Histones* class are shown in blue).

Finally, the functional enrichment of mouse IGs revealed biological pathways associated with genetic and protein regulatory processes including detection of chemical stimulus involved in sensory perception of the bitter taste, chromatin silencing, positive regulation of peptidyl-serine phosphorylation of STAT protein, and nucleosome positioning (−log10 −34.23 > −3.27). Other functions detected were immune, neuro-specific, and development processes such as mmu05322-Systemic lupus erythematosus, R-MMU-6805567 Keratinization, R-MMU-500792 GPCR ligand binding, R-MMU-1266695 Interleukin 7 signaling, hard palate development, and noradrenergic neuron differentiation (−log10 −29.94 > −3.75) (**Figure 3**).

**Figure 3.**
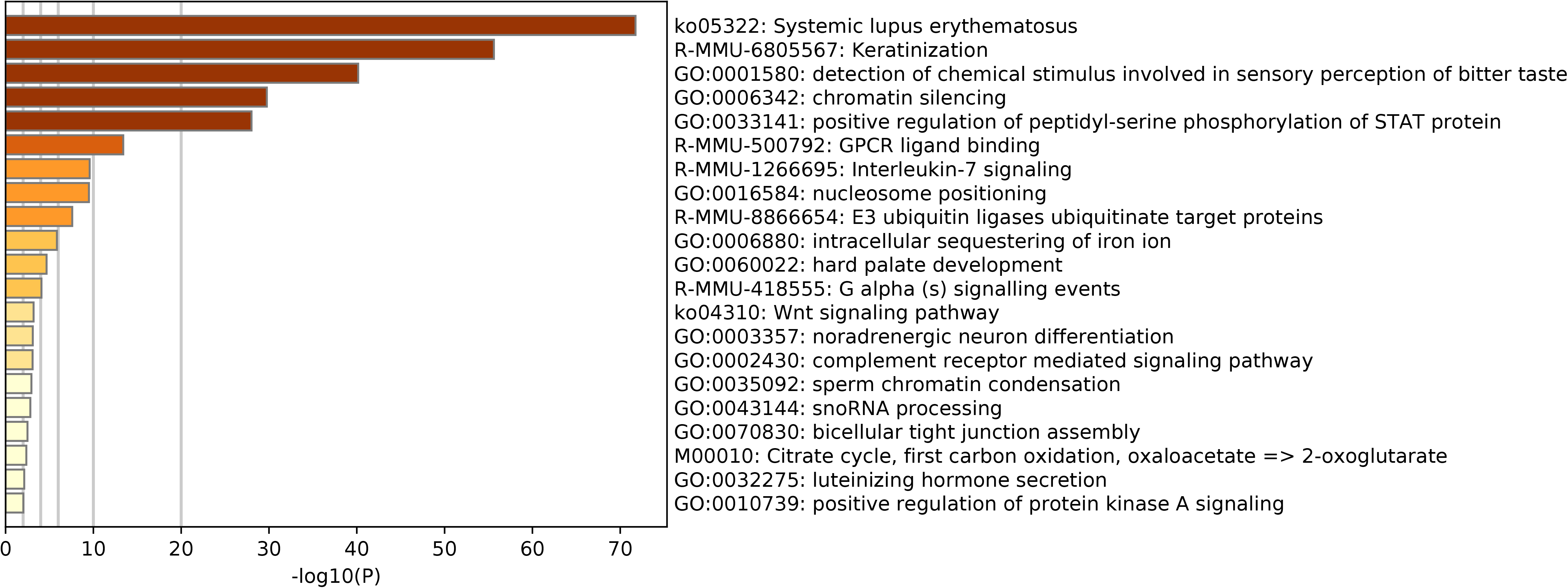
Functional enrichment of the mouse IG proteins. Gene ontology enrichment and pathways for IG proteins. The color key from yellow to brown indicates high to low *p*-values, respectively.

Altogether, functional assignment analysis suggests that IGs and MEGs proteins have distinctly and prevalent biological roles.

### Up-regulation of IGs reveal their regulatory role on neuro-specific functions across mouse development

Previous studies detected enrichment of neural-related functions among IGs (Grzybowska 2012). Moreover, since the expression of MEGs is modulated by the balance between the rate of transcription elongation and the alternative splicing of exons (Fong et al. 2014), we hypothesized that the natural absence of splicing on IG mRNAs could confer them differential regulatory roles in complex biological processes. Therefore, it was our interest to identify and analyze IGs that are expressed in mice during brain development. For that purpose, we analyzed expression data from the developing mouse telencephalon at stages in which its patterning is taking place (E9.5 and E10.5) (Dennis et al. 2016).

Overall, the expression of IGs was lower than that of MEGs (**Figure 4a**), which is consistent with previous *in silico* observations (Meena K Sakharkar et al. 2005). Out of 1,116 transcripts, differential expression analysis was performed for 1,087, with 37 of them (3.4%) showing up-regulation and nine down-regulation (0.82%) from gestational day E9.5 to E10.5. Moreover, 387 (35.63%) did not change expression, and 653 (60.12 %) were not expressed during the analyzed stages (**Figure 4b**). Meanwhile, among MEGs, 1247 were upregulated (6.13%), 789 were downregulated (3.88%), 13,198 had no expression changes (64.93%), and 5090 did not show expression (25.04%) (**Figure 4b**). It is noteworthy that an inverse expression pattern of genes with no expression changes (more MEGs that IGs) and those not expressed (more IGs that MEGs) was found in this comparison.

**Figure 4.**
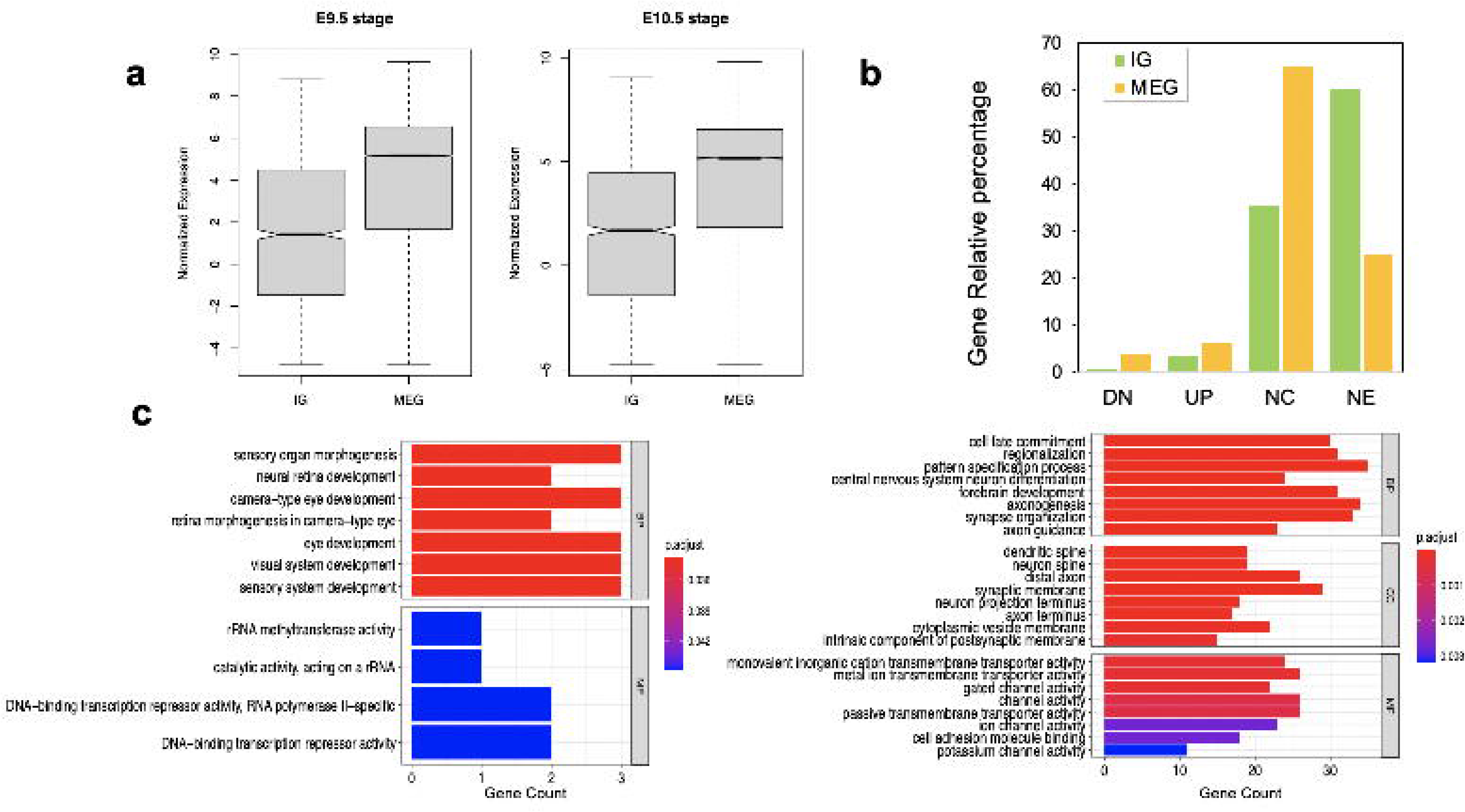
Expression levels of IGs in the mouse embryonic telencephalon compared to MEGs. **a)** IG and MEG normalized expression at 9.5 and 10.5 embryonic stages; **b)** IG and MEG, gene differential expression groups between 9.5 and 10.5 stages, (UP) upregulated, (DN) downregulated, (NC) not changes, (NE) not expressed; **c)** Significant enrichment of IG and MEG functions in upregulated genes, each barplot shows significantly enriched gene ontology (GO) terms on the y-axis, colored by their *p*-adjusted values, while gene count is represented on the x-axis. Test gene sets for enrichment analysis were the up-regulated genes in each dataset, and the background set was all up-regulated genes. GO terms are divided by BP (biological process), CC (cellular component), and MF (molecular function).

Our analysis revealed that all up-regulated IGs are exclusively enriched in biological pathways in eye and sensory organ development processes compared to MEGs also involved in other developmental and neural function pathways (**Figure 4c**). Moreover, significantly enriched terms in molecular function found for induced IGs are consistent with their regulatory role, including rRNA methyltransferase and DNA-binding transcription repressor activities. Opposite in the up-regulated MEGs the molecular function terms are highly enriched for transmembrane transporters and channel voltage activities (**Figure 4c**).

From the IG transmembrane protein group, transcripts for *Tram1l1, Cdk5r2, Nrarp, Kcnf1, Fzd7, Fzd8, Fzd10*, and *Cldn5,* were up-regulated. Strikingly, eleven members of the cluster of 22 *β-*protocadherins (pcdhbs) were among those up-regulated in the E10.5 telencephalon (nine IGs :*Pcdhb3, Pcdhb4, Pcdhb7, Pcdhb10, Pcdhb11, Pcdhb17, Pcdhb19, Pcdhb20, Pcdhb21,* and two SEGs *Pcdhb8 and Pcdhb22*)(**Figure 5a, Figure 6a**). Notably, our expression analysis additionally revealed up-regulation of *Olig1, Bhlhe22, Bhlhe23, Pou3f1, Pou3f2, Pou3f4, Foxq1,* and *Neurog1*, most of which are BHLH transcription factors crucial for the regulation of brain development and neuro-specific functions (**Figure 5b**). Moreover, regarding IGs within the histone group, *H2bc21, H2bu2, H2aw* were up-regulated during the mouse embryonic stages (**Figure 5c**). Finally, IGs from the others group with up-regulation observed are *Magel2*, *Magee1*,*Ndn* genes from the MAGE (melanoma-associated antigen) domain category, and *Pnmal2* (paraneoplastic Ma antigen family-like 2) from the PNMA category **(Figure 5d).**

**Figure 5.**
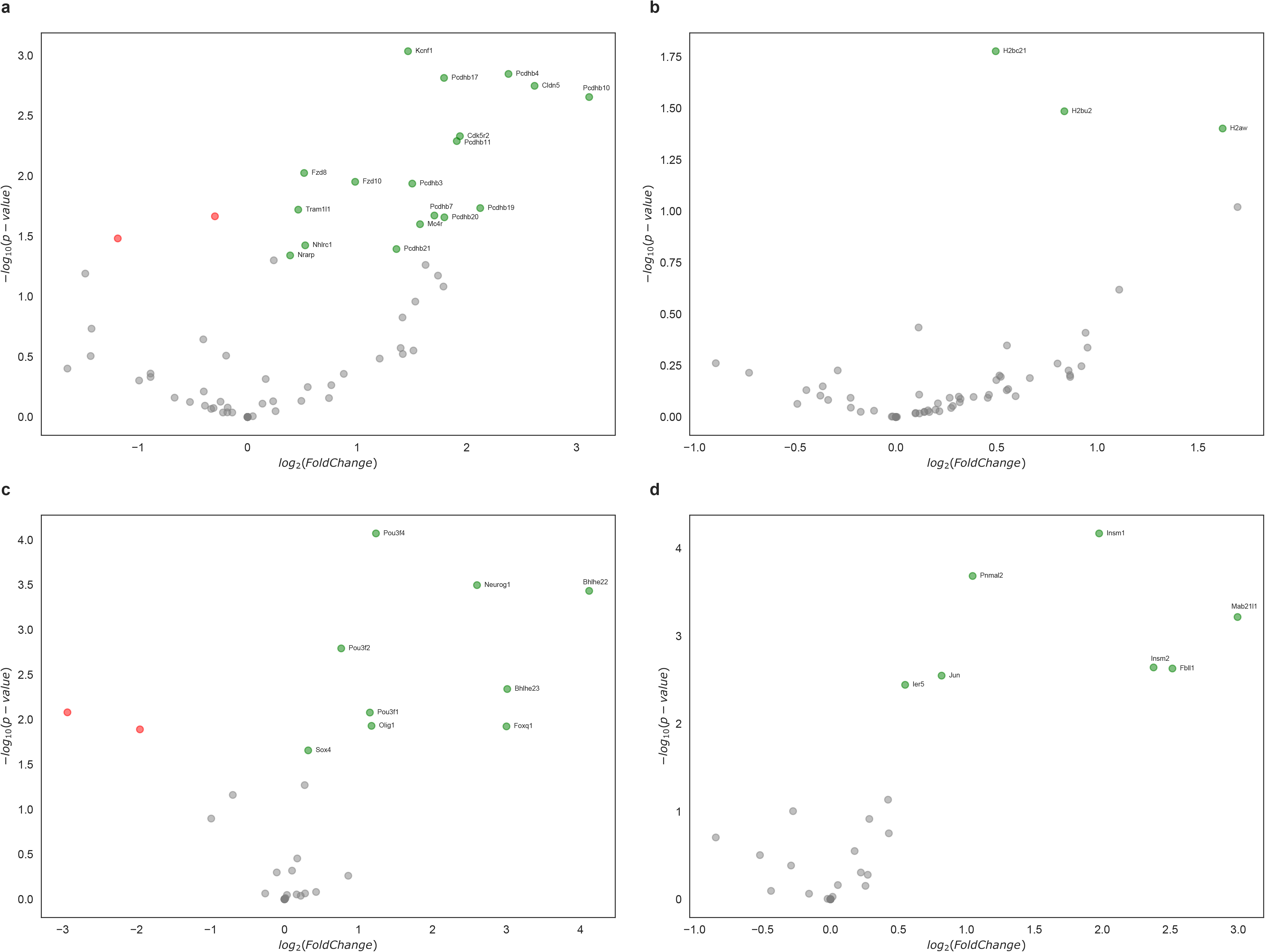
Differential expressed IGs in the mouse embryonic telencephalon. Expression of genes grouped by their Pfam assignment, determined by their log2 Fold change values. Up-regulated genes are highlighted in green, while downregulated genes are colored in red. **a)** Gene expression of transmembrane proteins, **b)** histones, **c)** transcription factors, and **d)** other protein families.

**Figure 6.**
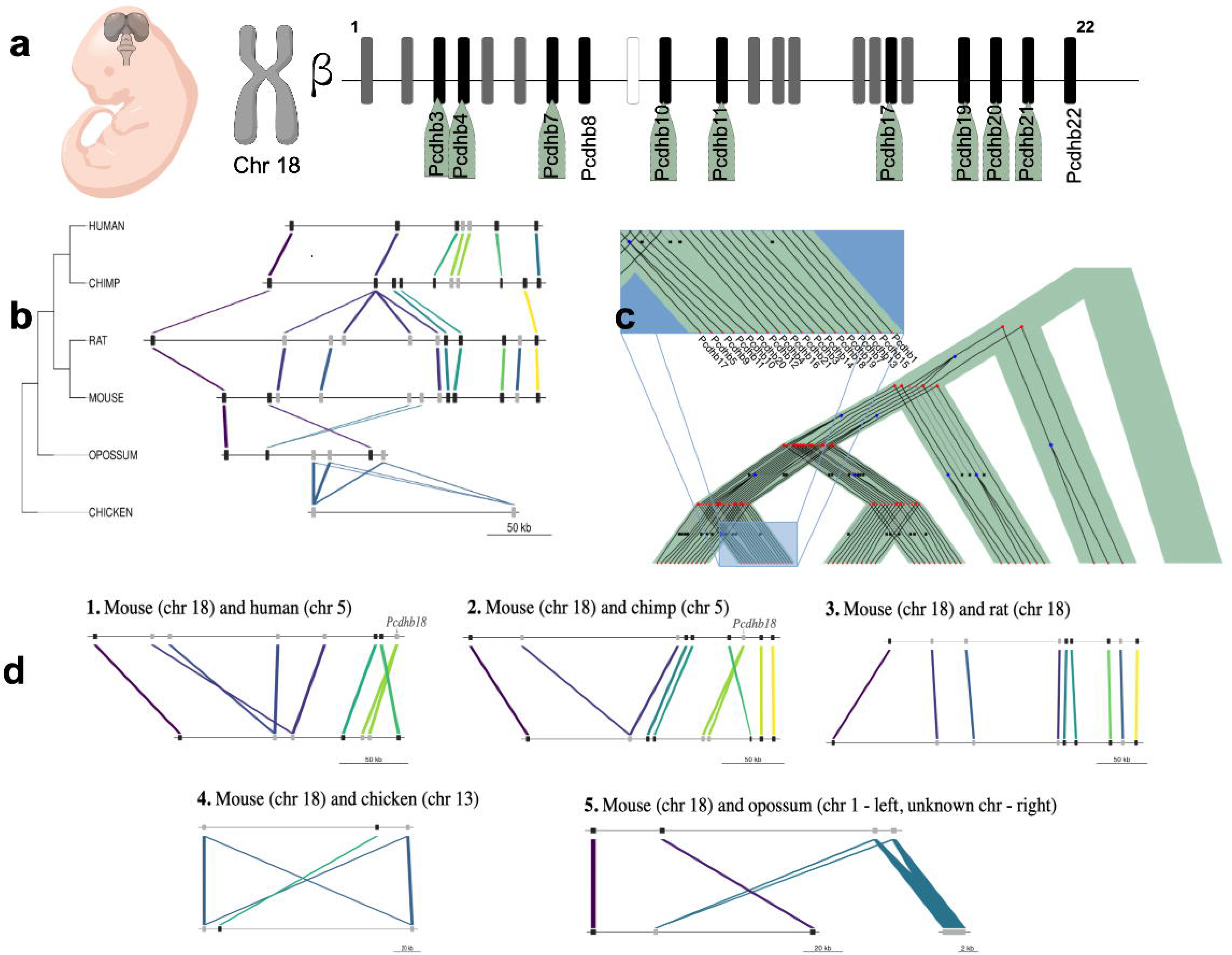
Expression and evolution analysis of the mouse cluster of β-protocadherins. **a)** Cluster of the 22 β-protocadherins in the mouse eighteen chromosome. Depicting in black the up-regulated genes in telencephalon tissue between mouse 9.5 and 10.5 embryonic stages, in gray up-regulated genes with no changes between stages, in white not expressed gene; **b)** Syntenic map of the ~300 kb β-protocadherin locus across five mammalian lineages and chicken as an outgroup. Colored lines depict orthology relationships across the phylogenetic tree. Genes connected by the same color belong to the same orthogroup identified by Proteinortho. Gene models are shown as black squares for single-copy orthologs and gray squares for expanded genes; Evolutionary events (blue squares represent duplications, red bullets represent speciations; **c)** Reconciliation tree of the protocadherin gene trees and the species tree. Each gene tree represents an orthogroup, internal nodes represent evolutionary events (blue squares represent duplications, red bullets represent speciations) and black crosses in leaves represent gene loss events; **d)** Pair-wise syntenic comparisons of the mouse β-protocadherin locus to four mammalian genomes and chick highlighting the lineage-specific loss and expansions of the *Pcdhb18* gene.

### The β-protocadherin gene cluster displays a high degree of syntenic conservation across mammalian genomes

To gain further insight on the evolutionary conservation of IGs with a functional role in telencephalon patterning, we studied the syntenic conservation of the ortholog genes of the 18 mouse IGs *β-* protocadherins (**Figure 6a**) in our set of seven species. We identified that human, chimp, rat, opossum, and chick contain copies of the mouse single exon *pcdhb* genes, while zebrafish do not. Overall, all the orthologous genes of the cluster are located in a single locus in their respective genomes, with varying lengths ranging from ~128kb to ~310kb, and displaying a few local inversions (**Figure 6b**). These results are consistent with previous studies that have explored the syntenic conservation of *pcdhb* genes across other vertebrate species (Noonan et al. 2004; W.-P. Yu et al. 2008). Even though we found syntenic conservation of some members of the β-protocadherin cluster, we observed slight disruptions of this order due to gene expansions, which can be either gene duplications or de novo formation. These gains are most notorious in the mammalian genomes (**Figure 6b**), suggesting gene expansion of the intronless *β-*protocadherins could be relevant for their role in neurogenesis, as well as other neuro-specific functions associated with the Wnt canonical pathway.

Then, by assessing syntenic conservation in a pair-wise fashion, we found relevant lineage-specific gene losses (**Figure 6c**). For instance, the *Pcdhb18* gene is absent in rats while it is present in mice and duplicated in primates. Although this gene is expressed in mice after the E10.5 stage their role in neuronal development has been reported (Bult et al. 2019). Together this evidence suggests that the nervous system complexity characteristic of mammalian species could be associated with the presence of single exon duplicated genes instead of splicing-derived protein isoforms as a neuron self-avoidance mechanism.

We looked into more detail at the evolutionary histories of *β-*protocadherins, by reconciling the gene trees of this gene cluster with the taxonomic species tree (**Figure 6d**). First, we observed that none of these genes is conserved in zebrafish. Moreover, some genes are gained in specific lineages, for example, *Pcdhb17* is only observed in mice and rats, while eight genes are shared across the mammals in the study. Three *β**-***protocadherins appear to be shared among primates and the marsupial opossum, while only *Pcdhb7, Pcdhb15* and *Pcdhb19* are shared between mouse and chick, and across other intermediate species, suggesting that these are the oldest *β-*protocadherins that give origin to the rest of them.

### Characterization and functional role of post-translational modifications in mouse IG proteins

In addition to alternative splicing, and mRNA editing, post-translational modifications (PTMs) constitute a defining factor of the complexity of proteomes by increasing structural and functional diversity of each proteoform, the set of multiple protein molecules encoded by one gene. Hence, protein PTMs have an essential role in protein structure-function features, including activity, stability, folding, and turnover (Uversky 2015). Since IGs fit the “one gene-one protein” concept, in which each gene encodes a single protein, we aimed to determine whether PTMs represent exclusive mechanisms of regulation for these genes. In our analysis, we observed that Succinylation 2.34 % and 2.27 % (for IGs and MEGs, respectively, in this and following paragraph), and S-nitrosylation (1.19 %, 1.53 %) were the PTMs with similar prevalence in both groups. These were followed with much lower frequency by Glutathionylation (0.89 %, 1.23 %), Glutarylation(0.26 %, 0.40%), Palmitoylation (0.18 %, 0.26 %) and Oxidation (0.07 %, 0.14 %)(**Figure 7a**). In contrast, PTMs with a unique presence in MEGs were Nitration (0.017 %), Myristoylation (0.01 %), Sulfation (0.0069 %), Pyrrolidone carboxylic acid, Carboxylation, and GPI-anchor (**Figure 7b**).

**Figure 7.**
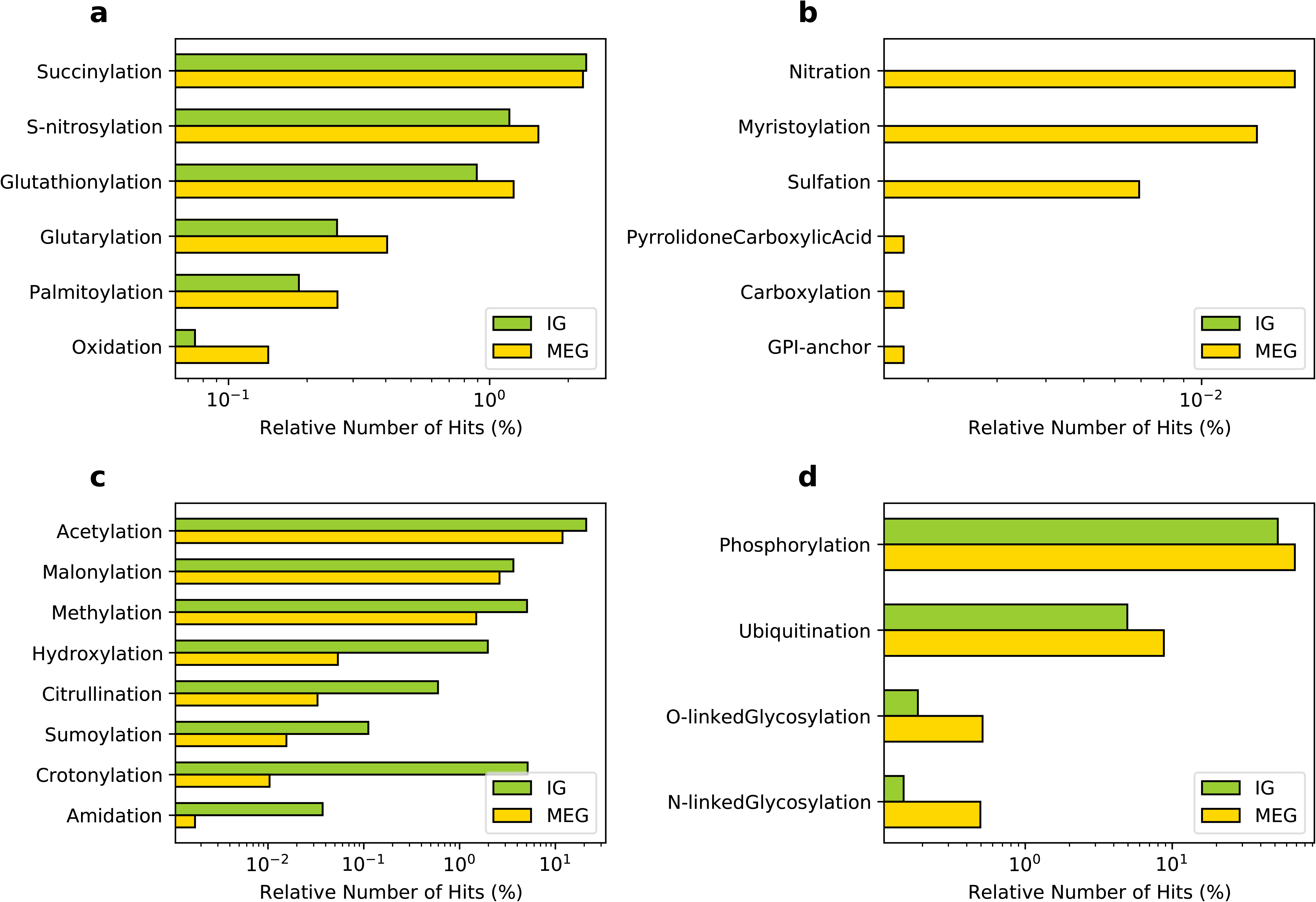
Distribution of post-translational modifications in the mouse IG compared to MEG proteins. **a)** PTMs with similar distribution between IG and MEG proteins. **b)** PTMs exclusive to MEGs. **c)** PTMs predominant in IG, and **d)** MEG proteins.

Notably, in accordance to their protein assignment, a differential enrichment for IG proteins was observed of Acetylation (20.98 %, 11.87 %), Crotonylation (5.13 %, 0.01 %), Methylation (5.05 %, 1.4 9%), Malonylation(3.64 %, 2.61 %), and Hydroxylation (1.97 %, 0.05 %) (**Figure 7c**) which are characteristic features of the core histones. Other PTMs of IG proteins, albeit at lower frequency, were Citrullination (0.59 %, 0.03 %), Sumoylation (0.11 %, 0.01 %) and Amidation (0.037 %, 0.0017 %)(**Figure 7c**). The complementary group of PTMs more enriched in MEG proteins included Phosphorylation (52.23 %, 68.23 %), followed by O-linked Glycosylation (0.18 %, 0.51 %), Ubiquitination (4.94 %, 8.76 %) and N-linked Glycosylation (0.14 %, 0.49 %)(**Figure 7d**). PTMs classified based on the modification-enabled functionality for membrane localization such as Myristoylation and GPI-anchor were found to be more frequent for MEGs (**Figure 7d**).

### Functional assignment of IGs as microproteins

The regulation of multidomain proteins at the post-translational level can be mediated by microproteins (miPs) (Staudt and Wenkel 2011) which are small proteins containing a single domain that form heterodimers with their targets and exert dominant-negative regulatory effects (de Klein et al. 2015; Eguen et al. 2015). In *Eukarya,* microproteins have been found to have a remarkable influence on diverse biological processes.

Aware of the differential occurrence of PTMs on IG proteins, the DNA binding repressor activity molecular function of up-regulated IGs during mouse brain development, and due to the remarkable role of miPs, we assessed whether this group of genes encoded proteins fitting the miP definition. Characteristic features of miPs are the short length of their primary structure, a homodimer domain, and negative modulating activity of protein multi-complexes. In the Pfam enrichment, we detected IG proteins with significance in the FYTT domain, which is a family of mammalian proteins of around 145-320 residues in length called 40-2-3 proteins with unknown function (http://pfam.xfam.org/family/PF07078). Our first approach was to analyze the peptide length of IG and MEG proteins. The highest length-frequency for IG peptides was in the range of 200-400 amino acids, compared to that of MEG peptides which was 300-500. Then, using the miPfinder tool (DOI: 10.1093/gbe/evx041), we identified the following microprotein IG candidates: the BHLH transcription factors *Bhlha9*, *Msg1*, *Ferd3l*, *Bhlhe23*, and *Ascl5* (e-value 4.6E-30), as well as the histones *H1f0, H1f1, Hils1, H1f2 H1f3, H1f4, H1f5, H1f6,* and *H1f10* (e-value 7.4E-09), corresponding to the H1 linker histone group.

### Conservation and Evolution of IGs across vertebrata

The origin of IGs has been explained by the mechanism of retrotransposition, which occurs by homologous recombination between the genomic copy of a gene and an intronless cDNA (Kaessmann, Vinckenbosch, and Long 2009). Retrotransposed single-exon genes also exist as pseudo-genes with lost molecular function, constituting almost half of the genes with one exon distributed in the mouse genome (**Suppl. Figure 2**). Protein coding IGs would have gradually evolved with novel molecular functions and the number and complexity of the gene repertoires of intron-containing and intronless genes are the results of subsequent duplication and deletion events throughout evolution.

To infer the evolutionary age of genes we implemented a bioinformatics method to assess the extent and patterns of distribution of each gene’s orthologs and paralogs in different species. The rationale of this approach is that widespread conservation of the orthologs of a gene in the different vertebrate taxa is an indication of old age for that particular gene. This approach allowed us to determine the conservation of IGs across 7 genomes, as well as to identify species-specific mouse IGs (**Figure 8**). In this analysis, we found that 543 out of the 1,116 mice IGs have orthologs in at least one of the other species. For the mammalian genomes, we found 442 genes conserved as IGs out of 501 orthologs in the rat genome, 335 orthologs in chimp with 250 conserved as IGs, 397 with 262 IGs in human, and 258 with 167 IGs in opossum (**Table 1**). Meanwhile, we found 133 orthologs in chick with 78 conserved as IGs, and 220 in zebrafish with 91 conserved as IGs (**Table 1**). We also identified out-paralogs of mouse IGs (genes that arose via duplication before a speciation) that are conserved in the other species: 36 in rat, 16 in human, 11 in chimp, 9 in opossum, 2 in chick and none in zebrafish. Finally, we identified 573 with no orthologs in the other species, suggesting that these are species-specific mouse IGs.

**Figure 8.**
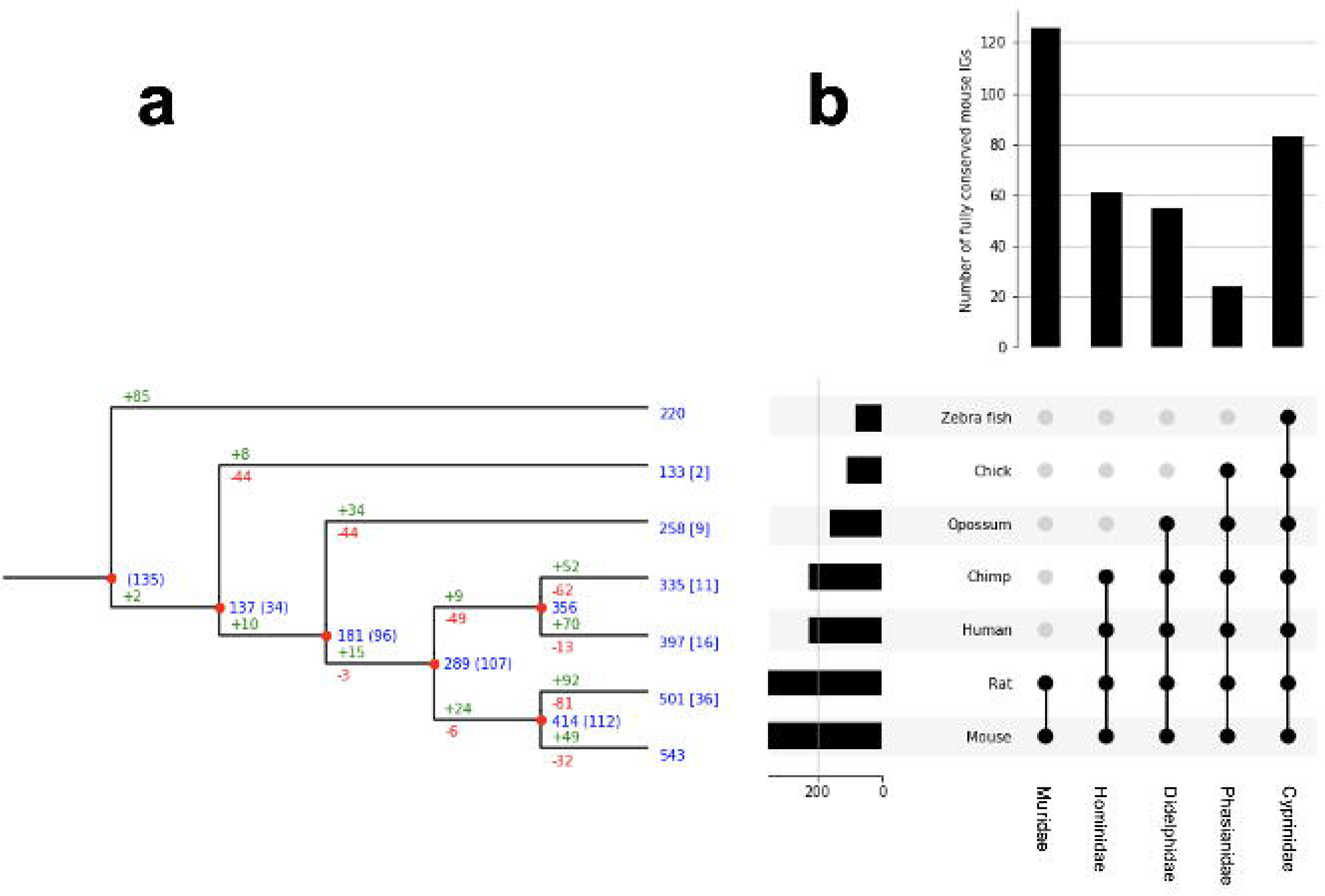
Clusterization of conserved mouse IG and MEG orthologs. **a)** Reconciliation tree representing the evolutionary history of mouse IG gene families. Blue numbers at each internal node represent the number of ancestral genes found in that clade that were inherited by an older ancestor. Blue numbers in brackets represent ancestral genes that might be generated at that evolutionary point, since they are not found in an outgroup of the clade. Green numbers represent gene gains due to duplications events and red numbers represent gene losses. Numbers at the leaves represent the number of orthologous genes in other species, and numbers in squared brackets represent the number of out-paralogs (genes that arose via duplication before speciation) in other species in the study; **b)** orthologous genes are grouped by age determined by the clade they are conserved in, the histogram on top of the upset plot shows the number of IGs that are specific to each clade, the histogram to the left of the upset plot represents the number of orthologous genes to mouse IGs for other species, and for mouse, the number of IGs for which an ortholog in another species was found.

**Table 1.** Summary of mouse IGs orthologs in selected genomes.

| Genome | IGs | uiSEGs | MEGs | Others | Total |
| --- | --- | --- | --- | --- | --- |
| Zebrafish | 91 | 0 | 90 | 39 | 220 |
| Chick | 78 | 1 | 54 | 0 | 133 |
| Opposum | 167 | 1 | 86 | 4 | 258 |
| Chimp | 250 | 0 | 59 | 26 | 335 |
| Human | 262 | 0 | 97 | 38 | 397 |
| Rat | 442 | 0 | 57 | 2 | 501 |

Moreover, in a parallel approach, we found out that 70% of the orthologous genes of mouse IGs, are IGs as well in the other genomes, only 2 are uiSEGs and 30% are MEGs, which supports the high conservation of the IG genetic structure across vertebra genomes (**Table 1**).

For comparative purposes, a similar analysis was performed for MEGs (**Suppl. Figure 4**). We found that for the mouse MEG dataset less than 5% of their orthologous genes are IGs and the rest are MEGs (**Table 2**).

**Table 2.** Summary of mouse MEGs orthologs in selected genomes.

| Genome | IGs | uiSEGs | MEGs | Others | Total |
| --- | --- | --- | --- | --- | --- |
| Zebrafish | 108 | 0 | 7602 | 1584 | 9294 |
| Chick | 122 | 2 | 7135 | 7 | 7266 |
| Human | 159 | 0 | 10304 | 966 | 11429 |
| Opposum | 523 | 8 | 8032 | 184 | 8747 |
| Chimp | 603 | 0 | 9520 | 374 | 10497 |
| Rat | 1171 | 0 | 10457 | 59 | 11687 |

From the previous analysis we clusterized IGs and MEGs into five age-groups named by the clade including the most recent common ancestor (MRCA) of each ortholog as inferred from the extant species analyzed. These groups were: *Cyprinidae*, *Phasianidae, Didelphidae, Hominidae, and Muridae* (**Figure 8b, Suppl. Figure 4b**). From the reconstruction of the evolutionary history of the mouse IGs, our results revealed that their conservation is more marked in the *Muridae* clade as it contains the largest number of orthologs common to its members (**Figure 8b)** followed in abundance by *Cyprinidae*. This indicates that a large number of IGs are sufficiently old to have orthologs in all *Cyprinidae* and that the clades that include the closest relatives to mice have increasing IG ortholog abundance. In contrast, the highest conservation of MEGs in gene numbers is among the *Cyprinidae* clade thereby revealing a much older age than that of *Muridae* IGs (**Suppl. Figure 4b**). For both IG and MEG orthologs, the number of paralog-related genes increases with age consistently with the rate of duplication of the edges of each clade (**Figure 8a**, **Suppl. Figure 4a**). Moreover, a significant number of in-paralogous genes in the zebrafish genome, generated via duplication after speciation, have an ortholog in the mouse genome.

With the purpose of determining whether there was a differential functional enrichment of IGs according to their evolutionary age, we analyzed the enrichment of molecular pathway GO terms in both IGs and MEGs. In agreement with a specialized role of IGs, our results show that IG and MEG orthologs are involved in different biological pathways although some shared pathways were detected as well (**Figure 9**). IG proteins with the MRCA conserved in *Cyprinidae* are histones highly enriched in negative regulation of megakaryocyte differentiation (−log10, −20.82). Other orthologs conserved to this group are linked in a lower level to thermogenesis, basal cell carcinoma, positive regulation of protein kinase A signaling, ribosomal large subunit assembly, wound healing, and vascular process in the circulatory system, platelet aggregation and development process such as cell-fate specification, negative regulation of animal organ morphogenesis, pituitary gland development, regulation of bicellular tight junction assembly(−log10, −9.33> −2.81). Meanwhile, gonad development is no enriched terms were found for orthologs exclusively shared with the *Phasianidae* ancestor. For *Didelphidae* we found G alpha signaling events (−log10, −8.86), while in the *Hominidae* group peptidyl-serine phosphorylation of STAT protein, and chromatin silencing was enriched (−log10, −6.47; −5.69). Noticeably, the most recent genes which belong to the *Muridae* group are exclusively enriched in intracellular sequestering of iron, complement receptor-mediated signaling pathway, and histone deubiquitination (−log10, −8.64>−3.63).

**Figure 9.**
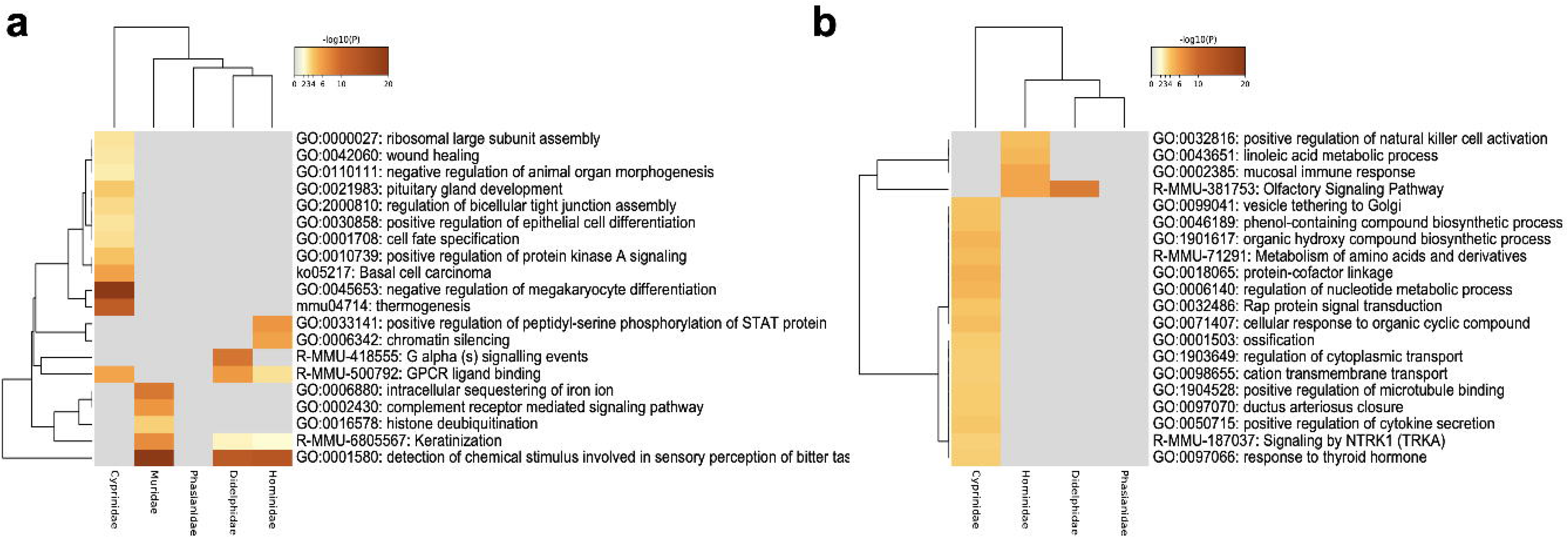
Functional enrichment of conserved mouse IG and MEG orthologs. **a)** Gene ontology enrichment and pathways for mouse IG conserved as IG in the selected species, orthologs are clustered regarding their “age” group. **b)** Gene ontology enrichment and pathways for a sample of 1,116 mouse MEG conserved as MEG in the selected species, orthologs are clustered regarding their “age” group. The color key from yellow to brown indicates high to low *p*-values, respectively.

Our analysis also identified IG proteins with enriched pathways shared among the various age groups. Detection of chemical stimulus involved in sensory perception of bitter taste, and keratinization are GOterms shared among *Muridae* (−log10, −22.91; −7.07)*, Hominidae* (−log10, −13.36;-2.10) and *Didelphidae* (−log10, −12.37; −2.39). Meanwhile, GPCR ligand binding is shared among *Cyprinidae* (−log10, −5.59)*, Didelphidae* (−log10, −6.11), and *Hominidae* (−log10, −3.05) (**Figure 9a**).

Then, we focused on determining the conservation of the biological role of IG orthologs among the different genomes. GO terms that are highly enriched in the seven genomes analyzed were detection of chemical stimulus involved in sensory perception of smell, organic substance metabolism, DNA packaging, signaling, multicellular organismal process, cell communication, transport, and localization, while GO molecular function terms enriched are olfactory receptor activity, Wnt protein binding, odorant binding, protein binding, catalytic and molecular transducer activity (**Figure 10**).

**Figure 10.**
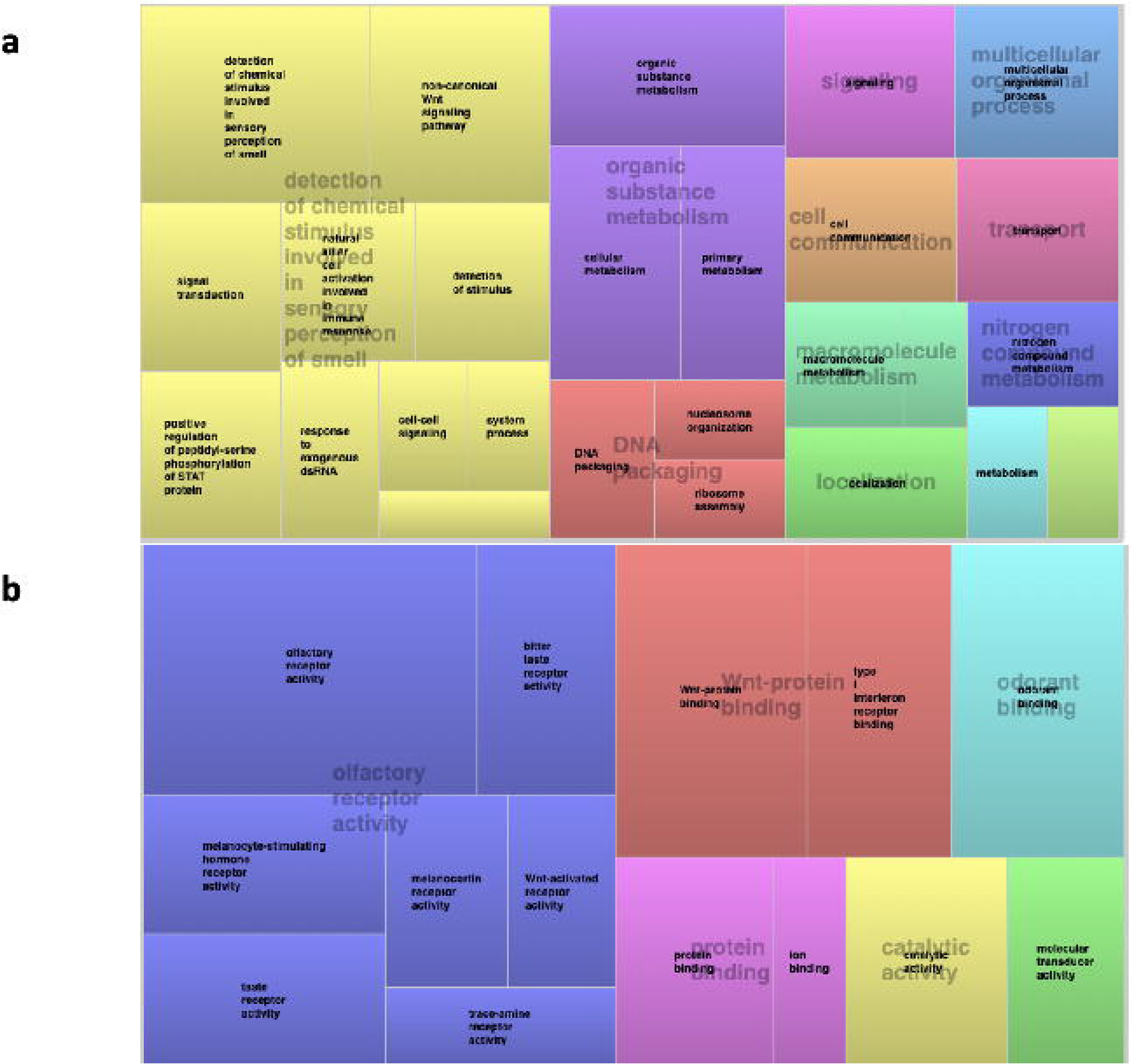
Conservation of the IGs functional role among orthologs. **a)** Overrepresented GO terms among all the conserved orthologs of mouse IGs for biological processes clusterized by their corresponding *p*-values; **b)** overrepresented GO terms among all the conserved orthologs of mouse IGs for molecular function clusterized by their corresponding *p*-values.

Finally, we studied the conservation of the genetic structure of the mouse orthologs in the other genomes. For that purpose, orthologs were clustered regarding their exon-intron architecture in those belonging to MEGs, SEGs, or IGs (**Table 1**, **Table 2**). This analysis revealed that most of the IGs identified in the mouse genome remained with this genetic structure in other species. As expected, due to its evolutionary closeness with the mouse, the genome with the highest conservation in gene architecture is the rat, with approximately 88% of conserved IGs. In dramatic contrast, the largest differences in gene structure were found for the zebrafish genome with 41% of conserved IGs. While the human genome is the one with the major presence of MEGs (**Table 1**). The transcript average determined for IG-MEG orthologs in all genomes was 2.23 transcripts per gene, being the human genome the one with the most abundant count of transcripts (up to 69 transcripts).

## Discussion

The mechanism of alternative splicing is a pivotal contributor to the diversity of proteins and the functional complexity of eukaryotic genomes. Intron-containing genes are capable of generating multiple protein isoforms by this process by which exons can be removed, lengthened, or shortened (Meena Kishore Sakharkar et al. 2005). The availability of detailed annotation of sequenced genomes for many organisms contributes toward a better understanding of their structure which has been shaped by flexible selection pressure. Studying the evolutionary dynamics of exon-intron patterns at the genomic level is likely to shed light on their role in genome structure and gene architecture. To deepen our insight into the structure and the evolution of mouse IGs, we examined their function, differential expression in the developing brain, the signatures for post-translational modifications of their encoded proteins, their potential as modulators of multiprotein complexes, as well as their evolutionary dynamics in comparison to their orthologues in other vertebrates.

### Functional assignment of IGs highlights prevalent and unique biological roles

The mouse genome annotated in the Ensembl database is composed of 22, 519 coding genes being those genes predominantly MEG type (80%) (https://www.ensembl.org/index.html). However, a considerable number of SEGs are present in this genome, and conservation of this fraction has been reported for other mammalian genomes (K. R. Sakharkar et al. 2006).

Our comparative analysis of the types of proteins encoded by IGs and MEGs revealed that these two populations have very divergent functional profiles. The most abundant types of proteins found among the former were the chromatin components histones and centromere proteins, transmembrane proteins of the G-protein coupled receptor family 1 and cadherins, and transcription factors containing BTB, forkhead, and HLH domains. In contrast, among MEGs, the most abundant proteins were those containing zinc finger, Pkinase, KRAB domain, and PH domains. These findings are consistent with previous findings that IGs are highly enriched in GPCRs and seven transmembrane domain proteins. These results reveal a functional divergence between IGs and MEGs which is likely to be the result of differential evolutionary constraints.

Among the transmembrane IG proteins, vomeronasal and taste receptors stand out as the most abundant. These proteins play a highly relevant role in chemoreception which is the most salient means of the interaction of *murinae* with their environment, with conspecifics, with potential prey or predators. These findings suggest that olfaction and taste receptors are required to be constantly transcribed in an efficient and rapid process, which could be the factor that favors their overrepresentation among IGs. Moreover, the taste receptor cells of vertebrates are continually renewed throughout the organism’s life which suggests a high demand for the housekeeping expression of these genes.

A considerable enrichment was also observed among IGs of BTB, Forkhead, HLH, HMG-Box transcription factors, and the chromatin components core histones H2AC, CENPTC, and the histone Linker cluster. An abundance of those IG proteins has DNA binding capacity. Thus, this indicates that IGs might be playing also an important role in chromatin assembly, packaging, and transcription.

Altogether, these results suggest that intronless genes have specialized roles, and a link to genetic expression regulation via chromatin remodeling. Overall, our results suggest that IG proteins have specialized, prevalent, and unique biological roles.

### Differential expression of IGs in telencephalon development, key genes for Wnt signaling

Embryonic development relies on the complex interplay of fundamental cellular processes, including proliferation, differentiation, and apoptosis. Regulation of these events is essential for the establishment of structures and organ development. The formation of the telencephalic architecture results from the interaction of the signaling centers located on the edges of the pallium. In this process, the Wnt signaling pathway plays an essential role in the dorsomedial pattern, where signals from the cortical hem direct the morphogenesis of the hippocampus, the corpus callosum and the generation of migratory Cajal-Retzius cells. From the lateral pallium, the anti-hem signals, EGF, FGF, Frizzled, and Sfrp determine the development of the olfactory cortex.

In a previous study, we highlighted the up-regulation of Wnt signaling genes of the canonical pathway in the early stages of the developing telencephalon in mice (Muley et al. 2020). The main receptors of the Wnt/beta-catenin signaling pathway are Frizzled domain proteins (Fzd), a family of seven-transmembrane G-protein coupled receptors that also possess a large extracellular cysteine-rich domain.

In this work, we found *Fzd7, Fzd8,* and *Fzd10* among the IG transmembrane proteins that were deregulated in the developing telencephalon, as well as a group of nine IGs and two SEGs of the protocadherin ß cluster. Previous studies have described *FZD8* as an essential receptor of the Wnt pathway implicated in brain development and size (Boyd et al. 2015). This gene is also highly expressed in two human cancer cell lines, indicating that it may play a role in tumorigenesis (Murillo-Garzón et al. 2018; Qiji Li et al. 2017). *FZD10* functions in the canonical Wnt/beta-catenin signaling pathway which may be involved in signal transduction during tissue morphogenesis (Z. Wang et al. 2005). In keeping with this, protocadherins have also been described as regulators in the Wnt signaling pathway (Mah and Weiner 2017). Moreover, protocadherins of the ß cluster, along with those of the α and γ clusters, act cooperatively in mice in olfactory-axon targeting, in the formation of diverse neural circuits, and in neuronal survival (Hasegawa et al. 2017; 2016). These functions, however, correspond to developmental stages that occur later than the one addressed in this study. Hence, our striking finding that half of the 22 ß protocadherins are up-regulated along with Wnt receptors during the development of the telencephalon, suggest that this group of mostly IGs has a differential function thus far unknown related to Wnt signaling at this early stage. Consistent with this idea, the Wnt binding molecular function GOterm is one of the most conserved among IGs in the genomes analyzed in this study.

Our expression analysis additionally revealed up-regulation of *Olig1, Bhlhe22, Bhlhe23, Pou3f1, Pou3f2, Pou3f4, Foxq1,* and *Neurog1*. Notably, this represents the up-regulation of three of the four members of the *Pou3* class of transcription factors. The POU genes encode a broad family of 6 classes (pou1f to pou6f) which are involved in developmental processes, mainly cell fate determination and differentiation (Tantin 2013). Among those, the four members of the pou3f class are preferentially expressed in ectodermal derivatives such as the developing mammalian nervous system (Bally-Cuif and Hammerschmidt 2003). The human *POU3F3* is an intronless gene also named *Brain-1*, which is a well-known transcription factor involved in the development of the central nervous system and its variant alleles have been associated with intellectual disability and language neurodevelopmental disorders (Blok et al. 2019). Furthermore, an important role of *NEUROG1* is as a promoter of proliferation or neuronal differentiation, while *Olig1* is involved in the generation and maturation of specific neural cells during the development of the spinal cord (Qi et al. 2016; Song et al. 2017). *Bhlhe22* and *Bhlhe23* in turn, are among those that were upregulated the most in mice during telencephalon development. In humans, *BHLHE22* has been identified as a highly methylated gene in endometrial cancer with potential epigenetic biomarkers in cervical scrapings (Liew et al. 2019), while Bhlhe23 has been linked to mammalian retinal development (Woods et al. 2018).

Finally, among the histone group, *H2bc21 (Hist2h2be), H2bu2 (Hist3h2ba), H2aw (Hist3h2a)* were also up-regulated. The H2B histone family members are responsible for the chromosomal fiber nucleosome structure in eukaryotes. *H2bc21/Hist2h2be* has been described in mouse as expressed in olfactory epithelium, while *H2bu2 (Hist3h2ba)* in neocortex and lens of camara type-eye, and *H2aw (Hist3h2a)* in retina (https://bgee.org/).In humans, *Hist2h2be* is a hub gene related to poor prognosis in rhabdomyosarcoma tumors in pediatrics patients (Qianru Li et al. 2019).

Summarizing, IGs appear to play crucial roles in the mouse telencephalon as deregulated genes are involved in gliogenesis, eye, and sensory organ development, canonical Wnt signaling, nucleosome organization, and have molecular regulatory roles. Therefore, in accordance with the functional assignment, our expression analysis supports that IGs play a critical role during mammalian brain development.

### IG proteins in the histone category have unique signatures and undergo specific PTMs

In accordance with the previously described finding of IGs specialized role linked to the regulation of gene expression via chromatin remodeling, we found unique and highly represented signatures in the histone protein category. Proteins encoded by mouse IGs have enriched signatures for histone H2A, and H2B, characteristic of key core histones involved in chromatin structure in eukaryotic cells, as well as linker histone H1/H5 and the CENB-type HTH. In addition to the identification of exclusive signatures, we compared potential regulatory mechanisms of IG and MEG PTMs. Although the variability of PTMs is high, these modifications are typically very specific and, altogether, 300 types are known to occur in proteins (Witze et al. 2007). Among all PTMs, we found that the most abundant (Phosphorylation and Acetylation) are differently represented among IG and MEG proteins.

As could be expected for IGs enrichment in chromatin remodeling protein domains, our results show that proteins encoded by IGs undergo specific PTMs for histone proteins such as crotonylation, methylation, sumoylation, citrullination, and sumoylation. However, our results suggest that IG-encoded histones have high specificity for Lysine-crotonylation, which is a recently identified post-translational modification associated with active promoters to directly stimulate transcription. Moreover, PTMs with changes in the physicochemical properties of amino acids like citrullination and amidation, are a characteristic feature observed highly enriched in IG proteins.

### Potential role of IGs as miPs in neural development and function

When we assessed the potential role of IGs as microproteins we found proteins with strong potential to be modulators of multi-protein complexes. Their targets or microproteins are mostly transcription factors that bind to DNA as dimers. In this study, we found potential miPs encoded by intronless genes that are bHLH transcription factors, with a regulatory role during critical events such as neural development and function. For example, *Ferd3l*, an evolutionarily conserved bHLH protein, is expressed in the developing central nervous system and functions as a transcriptional inhibitor. Another example is *BHLHe23,* a transcriptional regulator in the pancreas and brain that marks the dimesencephalic boundary (Bramblett et al. 2002), *Bhlha9* regulator of apical ectodermal ridge formation during limb development (Kataoka et al. 2018), *Msg1* which is predominantly expressed in nascent mesoderm, the heart tube, limb bud, and sclerotome during mouse embryogenesis (Dunwoodie, Rodriguez, and Beddington 1998), and *Ascl5* member of the ASCL family also referred as proneural genes which belong to a family of transcription factors that control the development of the nervous system, particularly neuroblast cell fate determination (Guillemot et al. 1993). *ASCL5* may be involved in the regulation of DNA-templated transcription. Moreover, its potential role in tumorigenesis has been described with upregulation in lung cancer and downregulation in brain tumors such as glioblastoma, anaplastic oligoastrocytoma, anaplastic oligodendroglioma, and oligodendroglioma (C. Wang et al. 2017). Additionally, consistent with the potential role in the development of IG-encoded miPs, we identified members of the H1 linker histone group that fit the criteria to be classified as miPs. These histone proteins belong to a complex family with distinct specificity for tissues, developmental stages, and organisms in which they are expressed (Izzo, Kamieniarz, and Schneider 2008).

### IGs and MEGs patterns of evolution in vertebrates

Analysis of the conservation of the orthologs of mouse IGs in seven species belonging to three classes of vertebrates revealed that the most numerous in each species were also IGs, with the exception of opossum. Individually, eutherians (rat, chimp, human) were the species with the most orthologs. Moreover, the clade that includes the mouse and its closest relative, the rat, had the largest number of conserved orthologs, this number decreases as the clades include gradually the more distantly related species and, strikingly, a distinctly larger number was found common to all species analyzed (clade Cyprinidae). While the IGs common to all species reveal the conservation of a population probably related to ancestral functions, those more abundant among species more closely related to mouse could be the combined result of the loss of IG orthologs during the divergence of the diverse vertebrate branches or of an increased rate of IG generation among mammals. Evidence in support of the latter possibility comes from our finding that in stark contrast, among MEGs, the most abundant group of orthologs corresponded to those common to all species and that the increase of orthologs among species more closely related to mice was not observed.

The two most abundant populations of IGs were found in cyprinidae and murinae. The first one contains the older IGs that are involved in diverse functions, among which nucleosome structure stands out; and the later which contains the more recent IGs which are involved in keratinization, GPCRs, and chemo detection by GPCRs in mammals. Moreover, the most ancient IGs code for histones that are conserved among all species, with some losses in opossum and chick. As histones are basic proteins known to be conserved across eukaryotes, it is not surprising that they are found to be some of the oldest IGs, since they play an important role in gene regulation. In some cases, histones are conserved as clusters, and in some others, these are generated in a specific lineage, due to multiple gene duplications.

Gene duplication is an important mechanism for the acquisition of new genes, frequently providing specialized or new gene functions (Magadum et al. 2013). Known mechanisms of gene duplication include retroposition, tandem duplication, and genome duplication (Pan and Zhang 2008). Our analysis shows that the vast majority (48%) of single exon genes in the mouse genome are a consequence of the retroposition mechanism. Moreover, regarding the duplication events, we found clear examples of IG tandem repeat cluster organization. Similar to the single exon beta gene cluster of protocadherins, we also observed that IG histones in the mouse genome are present as tandem families with the tendency to cluster in their chromosome organization. An example of this is the H2A histone family member L1J, a family of ten IG members in the mouse X chromosome, with only one ortholog (H2AL1RP gene) in human and one (histone H2A-beta) in opossum genomes. Almost all of the tandem repeat genes have parallel transcription orientation, which means they are encoded on the same strand.

The disrupted gene structure of most of the eukaryotic genes has led to a long-lasting debate regarding the origin of introns. The current state supports two alternative theories linked to the presence and absence of introns in primordial genes. Those are the “exon theory of genes” also known as “introns early”, and the “insertional theory of introns” or the “introns late theory” (Roy and Gilbert 2006). Another view is the synthetic theory of intron evolution which supports the hypothesis that merged both concepts (de Souza 2003). In accordance with the “introns early theory”, the comparison of mouse intron-bearing and intron-lacking IGs orthologs among the analyzed organisms, highlight that IGs might be more recent than MEGs.

The present study aimed to identify the conservation of the role of intronless genes in mammals and other vertebrate genomes. A comprehensive understanding of their biological function is essential to compare and contrast their evolution with that of intron-containing genes. Hence, we studied the complex regulatory role of intronless genes and their conservation in cellular environments using computational functional assignment, gene expression analysis, and evolutionary reconstruction.

First, we determined that the functions associated with IGs are very different from those associated with MEGs. Expression analysis of the developing telencephalon also revealed specific upregulation of IGs that encode genes involved in Wnt signaling, bHLH and POU transcription factors, and chromatin proteins. Among Wnt signaling-related proteins, it was striking to detect upregulation of half of all protocadherins of the ß cluster. Moreover, some IG transcription factors meet the criteria to be considered microproteins and thus appear to have modulated properties of protein complex formation. Overall, our results highlight a role for IGs as essential modulators of diverse biological processes as pivotal as cortical development, chemosensory functions, chromatin condensation, and gene silencing. In fact, specific modifications of IG proteins indicate that their regulatory roles extend to the post-translational level. Notably, some of the IGs highlighted in this study also have potential clinical relevance in humans. For example, *FZD8* and pcdhs associated with the Wnt signaling, an evolutionarily conserved regulatory pathway related to cellular and developmental processes such as cell fate determination and proliferation, which have also been identified as a key mechanism in cancer biology. Other IGs discussed in this study and linked with cancer and neurodevelopmental disorders were *POU3F3, BHLHE22, ASCL5,* and *Hist2h2be.*

Furthermore, the analysis of the evolutionary patterns of IGs revealed a large fraction of genes that appear to be of more recent generation as compared to the older and more conserved MEGs. Overall, this analysis reveals specific functions of IGs that distinguish them from MEGs and therefore strengthen the notion suggested by previous observations that these two groups are under differential evolutionary constraints.

## Author Contributions

K.A.P: project design, performed data collection, manuscript writing, proofreading, carried out bioinformatic analyses, prepared figures, and their interpretation. J.A.R.R: expertise in bioinformatic analysis methods, performed evolutionary reconstruction, prepared figures. G.E.H.O: literature search, bioinformatic analyses, and prepared figures. D. I. V: writing, bioinformatic analysis, and prepared figures. E. D. V: writing, bioinformatic analysis, and prepared figures. A.G.G: expertise in data analysis methods for the API-REST and data collection, prepared figures. V.M: proofreading, performed DEG analysis, prepared figures. A.V.-E: supervised the study and provided advice on the research strategy and manuscript writing. M.H.R: co-director of the study and project development, performed bioinformatic analysis and interpretation, writing, and proofreading.

## Acknowledgments

K.A.P acknowledges the CABANA program for training in bioinformatics. For technical support we thank Luis Alberto Aguilar Bautista, Alejandro de León Cuevas, Carlos Sair Flores Bautista and Jair García of the Laboratorio Nacional de Visualización Científica Avanzada (LAVIS). Critical comments and suggestions to this project development were received from Michael Jerzioski and Carlos Lozano Flores.

## Supplementary Material

The Supplementary Material for this article can be found online at:

**Code**: https://github.com/GEmilioHO/intronless_genes

**Supplementary Figure 1**. Bioinformatics pipeline for finding IG, uiSEG and MEG datasets

**Supplementary Figure 2.** Prevalence of intronless protein-coding genes among single-exon genes in the mouse genome.

**Supplementary Figure 3.** Enrichment of SUPERFAMILY assignments of mouse IG and MEG proteins.

**Supplementary Figure 1. Bioinformatics pipeline for finding IG, uiSEG, and MEG datasets.** Automatized steps (represented as 1-4): all protein-coding genes were retrieved for each species by accessing the Ensembl REST API platform. The number of exons for each gene were counted, and genes with two or more exons were identified as MEGs. Genes with one single exon were identified as SEGs and were subsequently filtered by their number of transcripts, keeping only those with one transcript. Using the IntronDB, we then classified these genes according to the presence of introns within their UTRs. Manual curation step (5): Genes with introns within these regions were identified as uiSEGs and those without, as IGs, while filtering out mitochondrial genes and genes with incomplete protein annotations. Output files are depicted as green squares.

**Supplementary Figure 2. Prevalence of intronless protein-coding genes among single-exon genes in the mouse genome.** All mouse genes having one exon are classified regarding their gene biotype, protein-coding intronless genes proportion is highlighted in pink.

**Supplementary Figure 3. Enrichment of SUPERFAMILY assignments of mouse IG and MEG proteins. a)** Enriched scop families in IG proteins: The scop families with the largest gene ratios are plotted in order of gene ratio. The size of the dots represent the number of genes in the significant background list associated with the scop family, while the color of the dots represents the adjusted p-values, **b)** Enriched scop families in MEG proteins: The scop families with the largest gene ratios are plotted in order of gene ratio. The size of the dots represents the number of genes in the significant background list associated with the scop family, while the color of the dots represents the *p*-adjusted values.

