## Supplementary figures and images for "Evolutionary Perspective and Expression Analysis of Intronless Genes Highlight the Conservation on Their Regulatory Role"

### Supplemental Figure 1.pdf

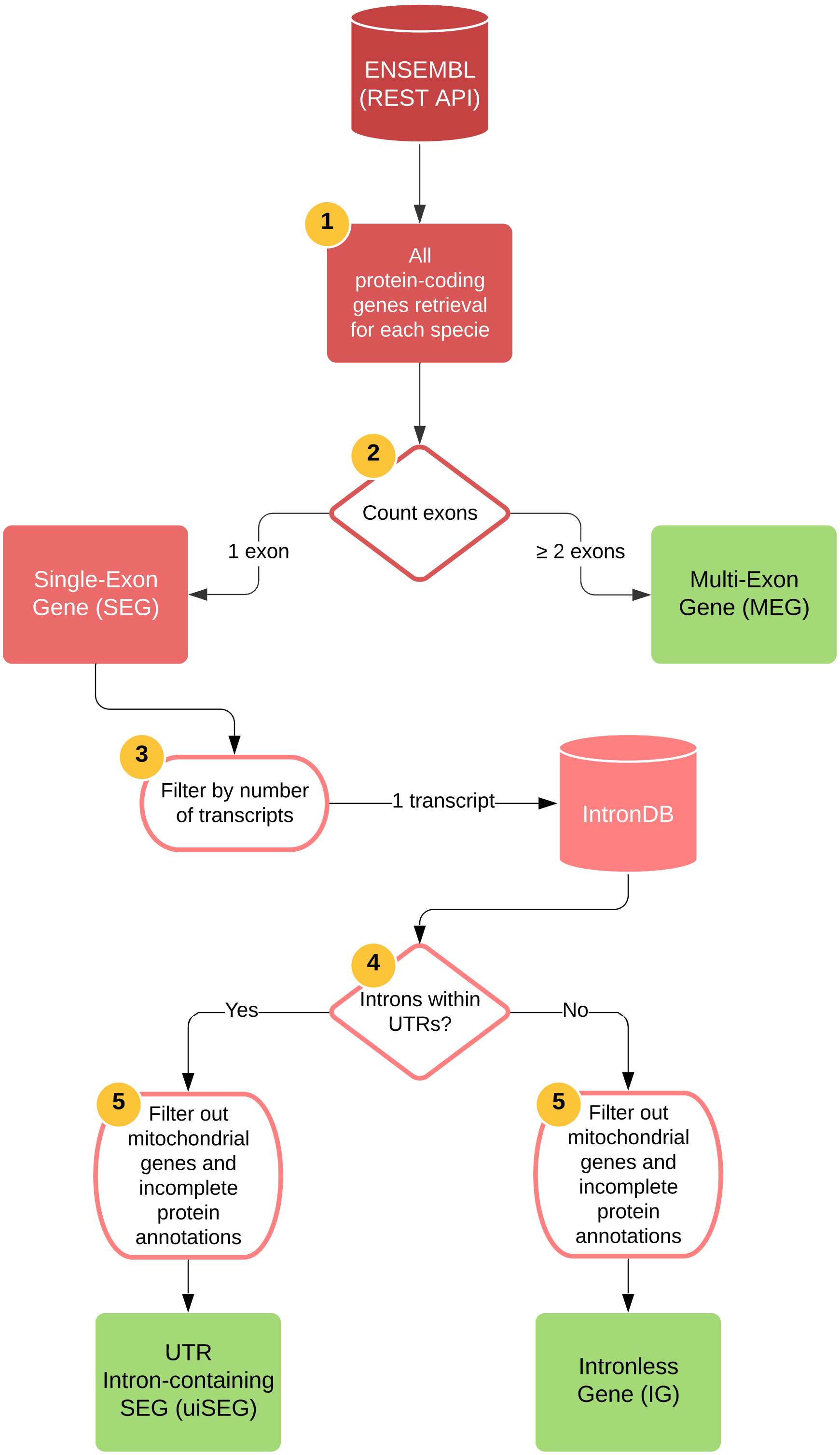

### Supplemental Figure 2.png

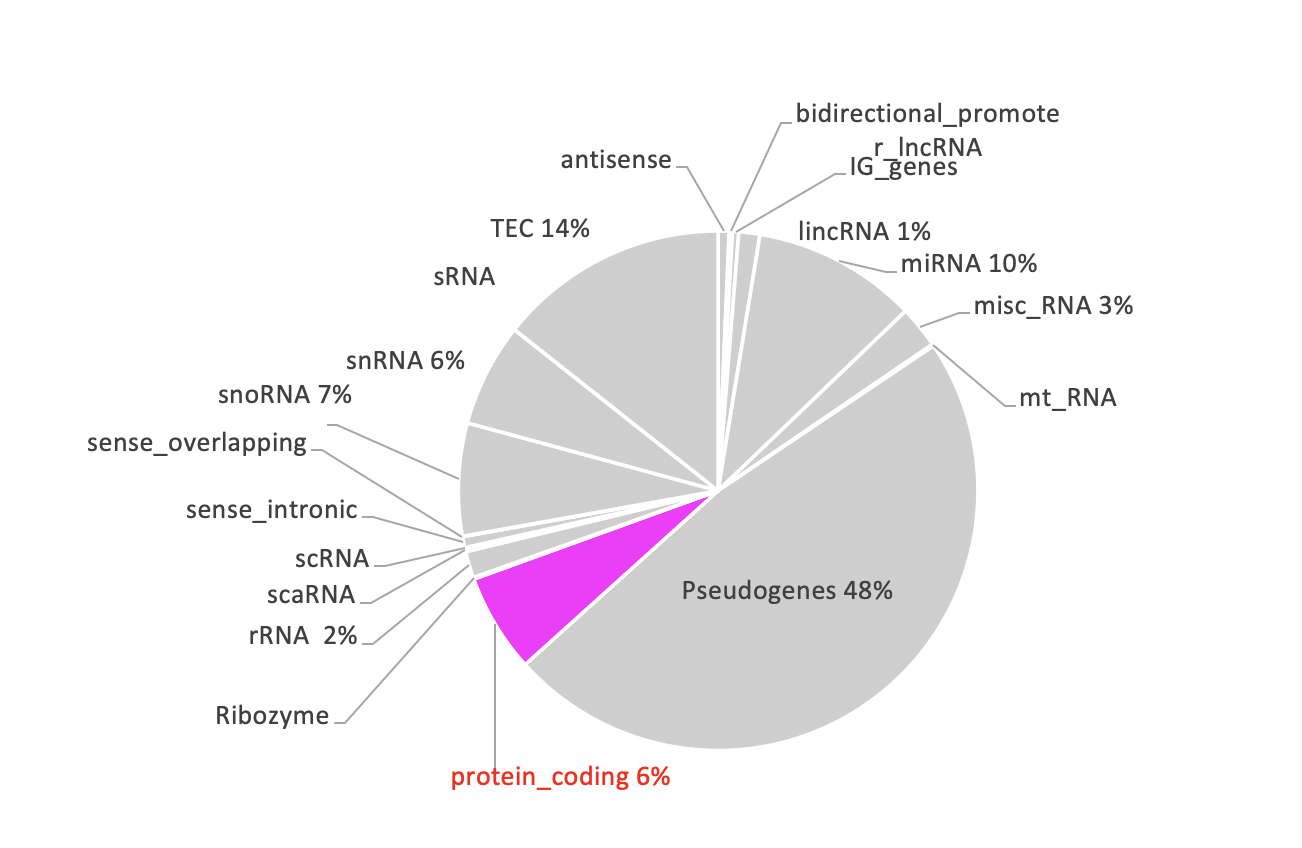

### Supplemental Figure 3.png

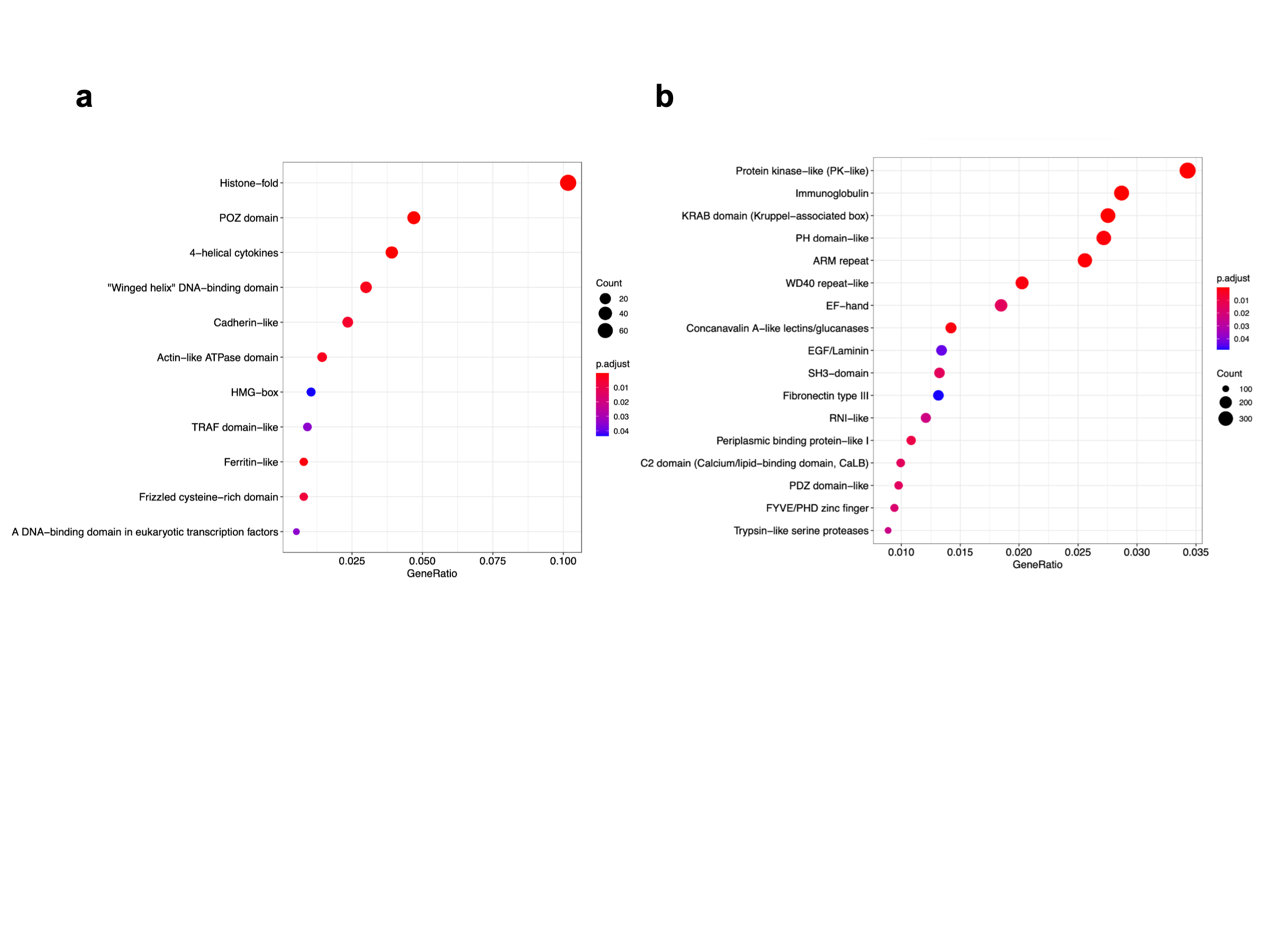

### Supplemental Figure 4.png

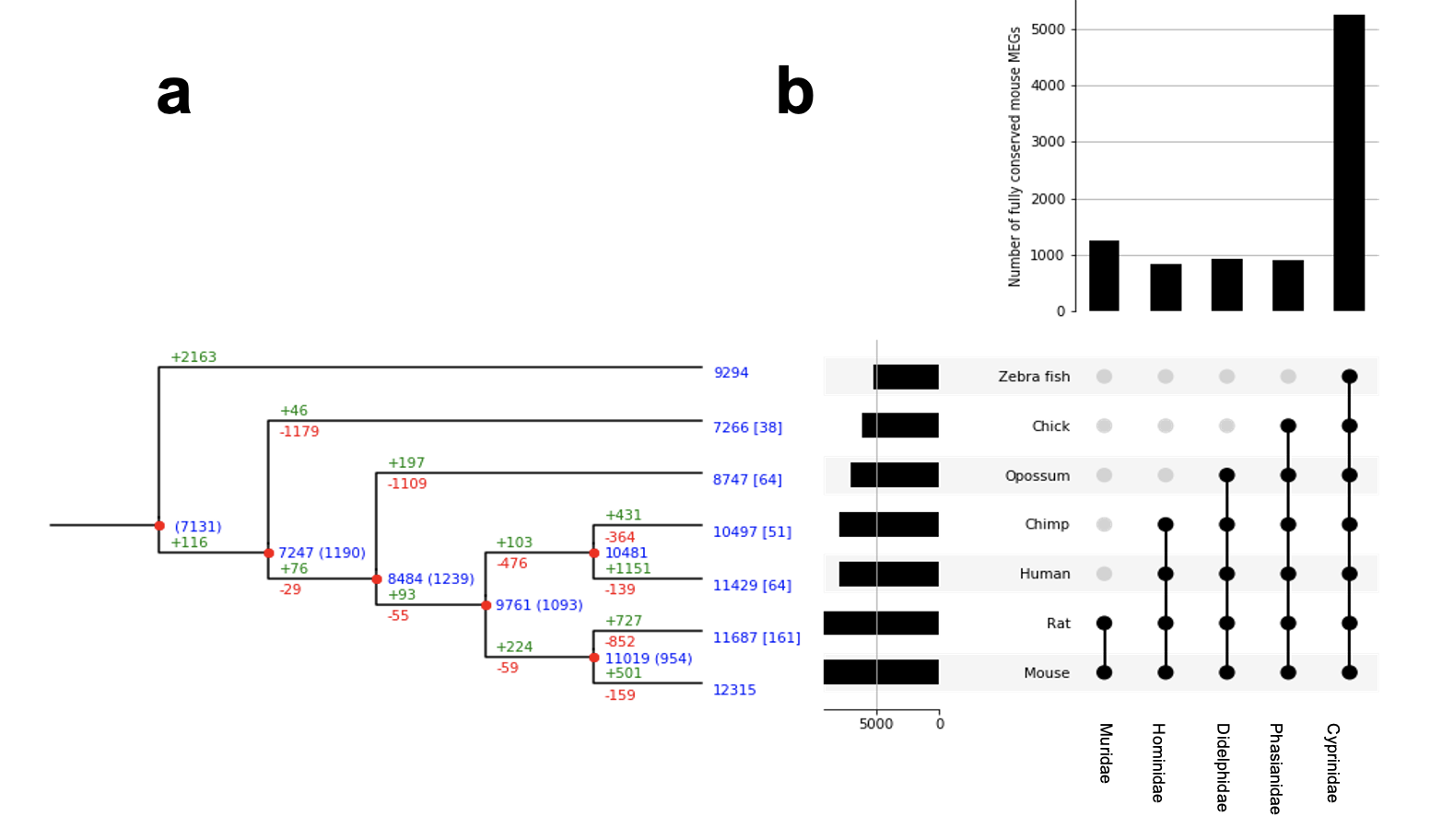
